# Learned adaptive multiphoton illumination microscopy

**DOI:** 10.1101/2020.08.14.251314

**Authors:** Henry Pinkard, Hratch Baghdassarian, Adriana Mujal, Ed Roberts, Kenneth H. Hu, Daniel Haim Friedman, Ivana Malenica, Taylor Shagam, Adam Fries, Kaitlin Corbin, Matthew F. Krummel, Laura Waller

**Affiliations:** Department of Electrical Engineering and Computer Sciences, University of California, Berkeley, CA, USA; Computational Biology Graduate Group, University of California, Berkeley, CA, USA; Berkeley Institute for Data Science; University of California San Francisco Bakar Computational Health Sciences Institute; Department of Pathology, University of California, San Francisco, San Francisco, California, USA; Department of Bioengineering, University of California, Berkeley, CA 94720, USA; Division of Biostatistics, University of California, Berkeley, CA 94720, USA

## Abstract

Multiphoton microscopy is a powerful technique for deep *in vivo* imaging in scattering samples. However, it requires precise, sample-dependent increases in excitation power with depth in order to maintain signal while minimizing photodamage. We show that cells with identical fluorescent labels imaged *in situ* can be used to train a physics-based machine learning model that solves this problem. After this training has been performed, the correct illumination power can be predicted and adaptively adjusted at each point in a 3D volume on subsequent samples as a function of the sample’s shape, without the need for specialized fluorescent labelling. We use this technique for *in vivo* imaging of immune responses in mouse lymph nodes following vaccination, with imaging volumes 2-3 orders of magnitude larger than previously reported. We achieve visualization of physiologically realistic numbers of antigen-specific T cells for the first time, and demonstrate changes in the global organization and motility of dendritic cell networks during the early stages of the immune response.

## Introduction

Imaging of cells *in vivo* is an essential tool for understanding the spatiotemporal dynamics that drive biological processes. For highly scattering tissues, multiphoton microscopy (MPM) is unique in its ability to image deep (200 *μ*m-2 mm, depending on the tissue) into intact samples. Because of the nonlinear relationship between excitation light power and fluorescence emission, scattered excitation light contributes negligibly to the detected fluorescence emission. Thus, localized fluorescent points can be imaged deep in a sample in spite of a large fraction of the excitation light scattering away from the focal point, by simply increasing the incident excitation power [1] (Fig. 1a).

**Figure 1:**
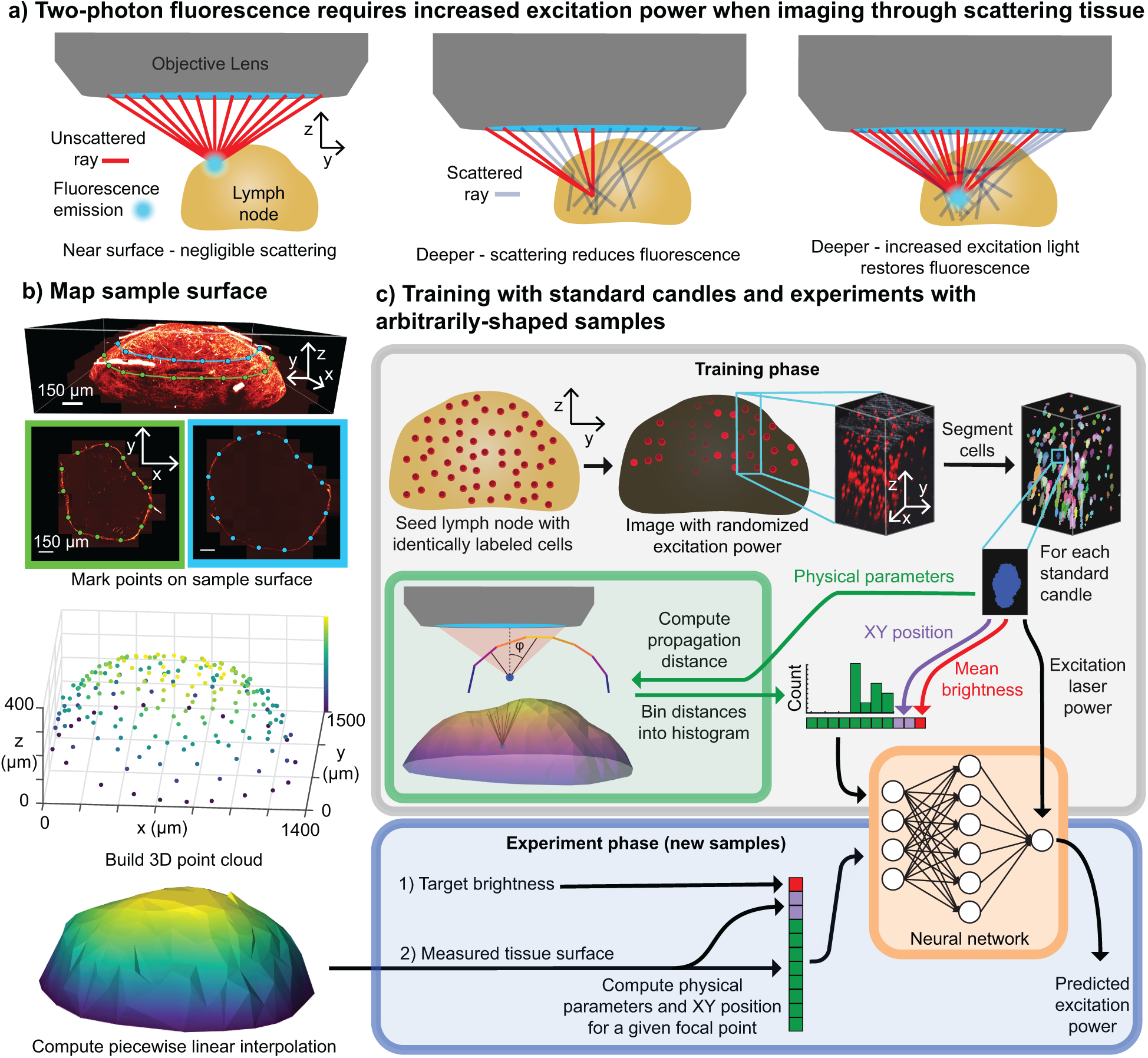
Learned Adaptive Multiphoton Illumination (LAMI). a) *In vivo* multi-photon microscopy requires increasing laser power with depth to compensate for the loss of fluorescence caused by excitation light being scattered or absorbed. b) The 3D sample surface is needed as input to our neural network. We map it by selecting points on XY image slices at different Z positions (top) to build up a 3D distribution of surface points (middle) that can be interpolated to estimate the sample shape. c) Training uses samples seeded with cells with the same fluorescent label (standard candles), which is imaged with a random amount of power. A 3D segmentation algorithm then isolates the voxels corresponding to each standard candle. The mean brightness of these voxels, position in XY field-of-view (FoV), and a set of physical parameters (a histogram of propagation distances through the tissue to the focal point at a specific angle of inclination to the optical axis (*ϕ*)) are concatenated into a single vector for each standard candle. The full set of these vectors is used to train a neural network that predicts excitation laser power. (Bottom) After training, subsequent samples need not be seeded with standard candles. The network automatically predicts point-wise excitation power as a function of the sample geometry and a user-specified target brightness.

Different points in the sample will incur different amounts of scattering and so determining the amount by which to increase the power for each position in 3D is a an important and nontrivial challenge, especially for larger samples that are not flat (e.g. large organoids, intact organs, large embryos). Too little power will fail to yield detectable fluorescence; too much will damage the structure being imaged. Taking the murine lymph node as an example, the total volume that can be imaged in a single experiment is often practically constrained to small subvolumes (10^−2^ mm^3^) over which the appropriate powers at each point in the sample are constant or predictable by a simple function [2]. Getting around this limitation requires a way to adapt excitation light power for each point in the sample individually while it is being imaged.

This practical constraint on imaging volume limits the range of biological investigations possible with MPM. For example, previous studies of T cell dynamics in intact lymph nodes have increased the density of transferred monoclonal T cells in order to achieve sufficient numbers for visualization (10^6^ or more) within the limited imaging volume of MPM. This number is three orders-of-magnitude more than the number of reported clonal T cell precursor (10^3^ − 10^4^) under physiological conditions [3, 4]. It is well-established that altering precursor frequencies changes the kinetics and outcome of the immune responses [5, 6, 7, 8], though it is not known how these alterations have biased the conclusions drawn by previous imaging studies.

Adaptive optics (AO) represents one strategy for addressing this challenge [9, 10]. By pre-compensating the shape of the incident excitation light wavefront based on the scattering properties of the tissue, the fraction of incident light that reaches the focal point increases, lessening the need to increase power with depth. However, AO still suffers from an exponential decay of fluorescence intensity with imaging depth when using constant excitation [1, 10], so an increase in incident power with depth is still necessary.

Alternatively, instead of minimizing scattering with AO, adaptive illumination (AI) techniques modulate excitation light intensity to ensure the correct amount reaches the focus. This strategy has been employed previously in both confocal [11] and multiphoton [12] microscopy. The basic principle is to implement a feedback circuit between the microscope’s detector and excitation modulation, such that excitation light power is turned off at each pixel once a sufficient number of photons have been detected. However, a major limitation of this approach is that it doesn’t account for fluorophore brightness and labelling density; thus, it is impossible to disambiguate weak fluorophores (e.g. an autofluor) receiving a high dose of incident power from strong fluorophores (e.g. a fluorescent dye) receiving a low dose. In order to prevent an unlabelled or weakly fluorescent part of the sample from being overloaded with excitation light, a user-specified upper bound is set on maximum power, which can vary spatially by over an order-of-magnitude for highly-scattering thick samples. Thus, applying this approach to image 100s of *μm*s deep in such samples still requires additional prior knowledge about the attenuation of fluorescence in different parts of a sample. For a flat sample (e.g. imaging into brain tissue through a cranial window) this prior knowledge may be as simple as an exponential increase with depth. For curved samples, the function is more complex. As a result, imaging large volumes of highly-scattering tissue for non-flat samples has remained a challenging task.

Here, we provide a solution to this problem. Building on the idea of adaptive illumination, we utilize a combination of machine learning and computational modeling of optical physics in order to learn to the relationship between fluorescence intensity of a standard sample and incident excitation power, given the sample’s shape in a one-time calibration experiment. On subsequent experiments, this enables pre-compensation of the power of incident excitation light as a focal spot is scanned through each point in the sample in order to achieve sufficient numbers of detected photons throughout the volume, without the need for any specialized sample preparation. We describe a simple hardware modification to an existing multiphoton microscope (costing *<*$50) to enable the modulation of laser power as the excitation light is scanned throughout the sample. We call our technique learned adaptive multiphoton illumination (LAMI).

Our central insight is inspired by the idea of “standard candles” in astronomy [13], where the fact that an object’s brightness is known *a priori* allows its distance to Earth to be inferred based on its apparent brightness. Analogously, we hypothesize that by measuring the fluorescence emission of identically-labelled cells (“standard candles”) at different points in a sample volume under different illumination conditions, we could use a neural network to learn an appropriate adaptive illumination function that could predicted from sample shape alone.

Applying LAMI to intravital imaging of the murine lymph node, we first show that this function can be learned and that it generalizes across differently-shaped samples of the same tissue type (e.g. one murine lymph node to another - though we note that moving to a new tissue type, which would attenuate light differently, would require a new calibration experiment). Having learned the function in a one-time calibration experiment, the trained neural network can be used to automatically modulate excitation power to the appropriate level at each point in the sample, enabling dynamic imaging of the immune system with single-cell resolution across volumes of tissue two or more orders-of-magnitude larger than previously described. Unlike previous studies which artificially increased the number of monoclonal T cells to 10^6^ (2 orders of magnitude greater than typical physiological conditions) in order to visualize them in a small imaging volume, we image physiologically realistic (5 *×* 10^4^ transferred) cell frequencies for the first time.

## Results

### Learning illumination power as a function of shape

The detected fluorescence intensity at a given point results from a combination of: 1) The sample-dependent physics of light propagation (e.g. scattering potential of the tissue, fraction of emitted photons that are detected, etc.), factors that are difficult to model*a priori* due to heterogeneity in sample shapes. 2) The fluorescent labelling (e.g. the type and local concentration of fluorophores), a nuisance factor that makes it difficult to disambiguate weak fluorophores receiving a high dose of incident power from strong fluorophores receiving a low dose.

Our method relies on the fact that, if fluorescence labelling of distinct parts of the sample are, on average, constant (i.e. “standard candles”), we can separate out the effects of fluorescence strength and tissue-dependent physics by performing a one-time calibration to learn the effect of *only* the tissue-dependent physics for a given tissue type. This calibration captures the physics relating excitation power, detected fluorescence, local sample curvature, and position in the XY field-of-view (FoV), which includes optical vignetting effects. By generating a dataset consisting of points with random distributions over these variables, we can train a neural network to predict excitation power as a function of detected fluorescence, sample shape, and position. On subsequent experiments in different samples of the same type, the excitation power required to achieve a desired level of detected fluorescence can be automatically predicted for each point in the sample based only on sample shape and XY position.

In the mouse lymph node, we can easily introduce standard candles by transferring genetically identical, identically-labelled (with either cytosolic fluorescent protein or dye) lymphocytes, which then migrate into lymph nodes and position themselves throughout its volume. Although there are certainly stochastic differences in labeling density between individual cells, as long as these differences are not correlated with the cells’ spatial locations, they will not bias the calibration.

The sample shape must be measured in such a way that: 1) it effectively captures the relevant physics of fluorescence attenuation, and 2) It can be used as input to a neural network. First, points were selected on the sample surface in XY images of a focal stack to generate a set of 3D points (Fig. 1b). These points were interpolated in 3D in a piece-wise linear fashion to create a 3D mesh of the sample surface.

Next, we used this mesh to compute statistics that captured important properties of the physics of the imaging process. In multiphoton microscopy, the distance light travels through tissue is an important quantity, as both the fraction of excitation light that attenuates from scattering/absorption and the fraction of fluorescence emission that absorbs are proportional to the negative exponential of this distance [1], assuming homogeneous scattering. We thus reasoned that measuring the full distribution of path lengths (i.e. every ray within the objective’s numerical aperture), would provide an informative parameterization to predict fluorescence attenuation. Empirically, we found that the full distribution of distances was not needed to achieve optimal predictive performance, and that measuring twelve distances along lines with a single angle of inclination relative to the optical axis was sufficient (**Fig. 1c**, green box). Since we expect our optical system to have rotational symmetry about the optical axis, the measured distances were binned into a histogram, and the counts of this histogram were used in the feature vector fed into the neural network.

Our neural network takes inputs of mean standard candle brightness, local sample shape, and position within the XY FoV and outputs a predicted excitation power (Fig 1c, orange box). Next, we need to generate an appropriate training dataset. Each example in our training dataset is a single standard candle cell that was imaged with a random, known amount of excitation power. Neural networks are excellent interpolators and poor extrapolators, so we ensured that the random excitation power used in training induced a range of brightnesses spanning too-dim-to-see to unnecessarily bright (Fig. 1c, top middle, **Movie S1**). Unlike contemporary deep neural networks [14], the prediction only requires a very small network with a single hidden layer. Once trained, the network can then be used with new samples to predict the point-wise excitation power needed for a given level of brightness (Fig 1c, bottom). After the one-time network training with standard candles, experiments can be fluorescently labeled without standard candles, and only the sample shape is needed to predict excitation power.

### Modulating excitation light across field-of-view

The appropriate excitation power often varied substantially across a single 220*x*220*μm* FoV, especially when imaging curved edges of the lymph node where the sample was highly inclined relative to the optical axis, thereby including both superficial and deep areas of the lymph node (**Fig. S1**). The trained network predicted very different excitation powers from one corner of the FoV to another in such cases. In order to be able to deliver the correct amount of power, we need to be able to spatially pattern excitation light at different points within a single FoV. To accomplish this, we designed a spatial light modulator (SLM) capable of modulating excitation laser power within the raster scan of a single FoV (**Fig. S1, S2**). By modulating excitation power over time, we could effectively achieve spatially-patterned excitation. The SLM was built using an Arduino-like programmable micro-controller and a small op-amp circuit, allowing it to retrofit an existing multiphoton microscope for less than $50 (**Supplementary materials and methods**).

### Generalization across samples

To validate the performance of LAMI and demonstrate that it can generalize across samples, we trained the network on one type of lymph node and tested on a differently-shaped lymph node from a different anatomical location (Fig. 2a). The experimental test lymph node was seeded with a variety of fluorescent labels and imaged *ex vivo* to eliminate the possibility of motion artifacts associated with intravital microscopy. The surface of the test lymph node was mapped as described previously (Fig. 1b). Several different desired brightness levels were tested to find one with appropriate signal. For comparison, we imaged the test lymph node with our method, with a constant excitation power, and with an excitation power predicted by a ray optics model (**Supplementary materials and methods, Fig. S3**). With constant excitation, fluorescence intensity rapidly decayed after the first 50 *μ*m. Adaptive excitation provided clear visualization of cells throughout the volume of the lymph node (Fig. 2b, **Movie S2**), up to the depth limit imposed by the maximum power of the excitation laser on our system of around 350 *μ*m (**Fig. 2e**). Though the ray optics model enabled similar depth, its predictions failed to yield detectatble fluorescence in areas where cells were clearly visible with adaptive excitation (**Fig. S4**).

**Figure 2:**
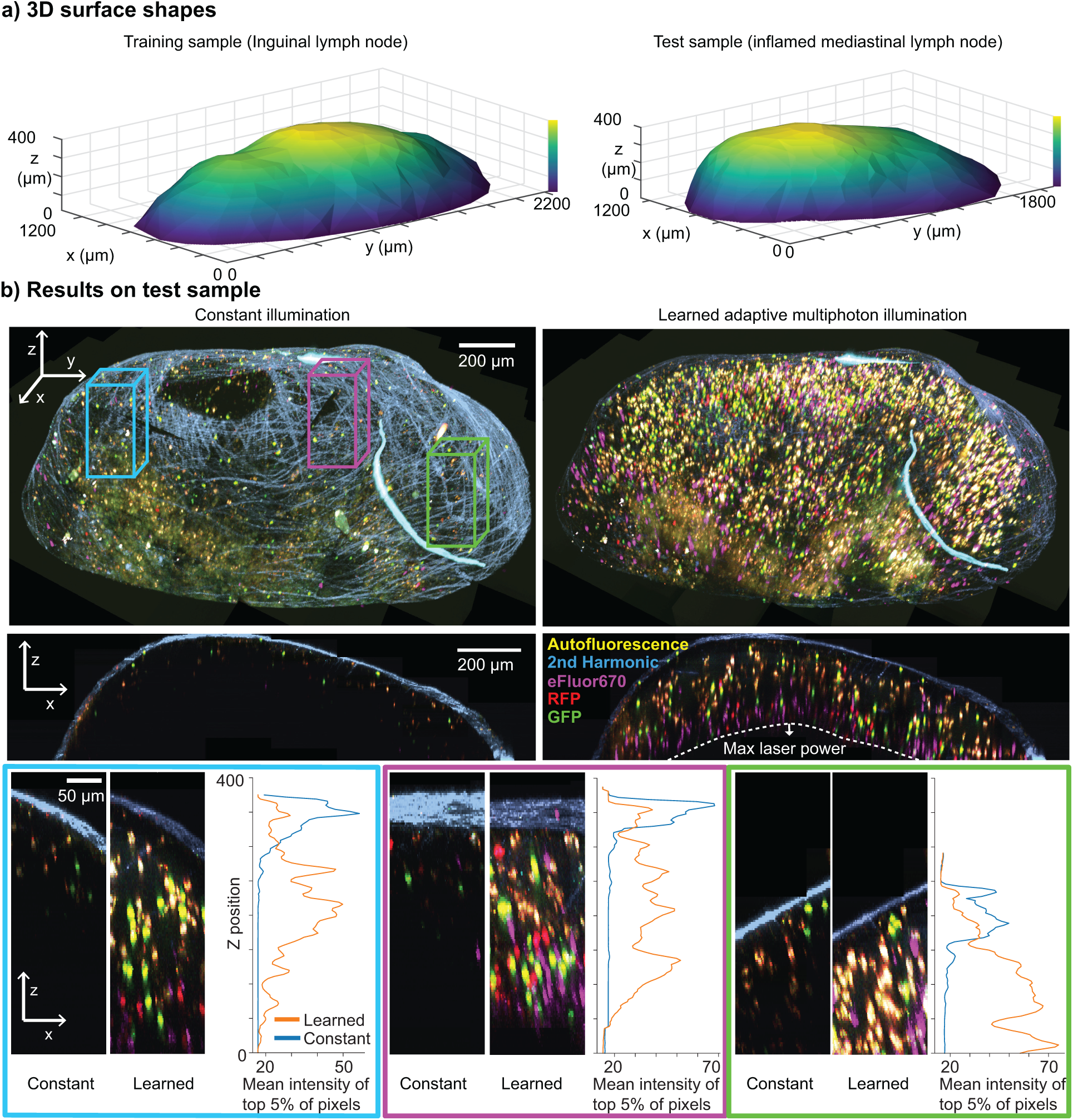
Validation of LAMI on lymph node samples. a) The surface shapes of lymph nodes used for (left) training with standard candles (inguinal lymph node) and (right) testing (inflamed mediastinal lymph node). b) Results with constant illumination power vs adaptive illumination power in the test sample, which had been seeded with lymphocytes labelled with GFP (green), RFP (red) and eFluor670 (magenta). Unlike constant illumination, which has very poor contrast at deeper depths, LAMI gives good signal throughout the imaging volume up to the maximum excitation laser power. (Top) 3D views, (middle) XZ projections, (bottom) XZ projections of representative areas with different surface curvature. Plots show Z-position vs mean intensity of top 5% of pixels to demonstrate good signal is maintained with depth using LAMI.

### Motion artifact correction and cell tracking

In order to conduct *in vivo* investigations, we must first address motion artifacts, which are an inescapable feature of intravital imaging and can compound when imaging large volumes over time. We thus develop a correction pipeline based on iterative optimization of maximum a posteriori (MAP) estimates of translations for image registration and stitching (**Fig. S5, S6, Supplementary methods**). We identified three main categories of motion artifacts (Fig S6b): 1) transverse (i.e. XY) movements due to breathing, 2) global XYZ drift of each Z stack over time, and 3) XYZ misalignment between different Z stacks. These corrections enabled the recovery of stabilized timelapses in which cell movements can be clearly visualized and tracked (**Movie S3, S4**).

Finally, we developed a machine learning pipeline for tracking cell locations over time based on their fluorescent labels (**Fig. S7**). Automating this process was essential, as each dataset contained up to thousands of labelled cells in up to twenty time points. Our pipeline enabled their detection across all datasets with the manual classification of no more than 500 cells for each time, a task that could be completed in a few hours of manual effort. Briefly, this pipeline consisted of two stages: a 3D segmentation algorithm to identify cell candidates, followed by a neural network that used hand-designed features (**Fig. S8**) to classify each candidate as positive or negative for a given fluorescent tag. We used an active learning [15] framework to efficiently label training data for this classification netowrk, which led to a 40*×* increase in the efficiency of data labeling (**Fig. S6**).

### *In vivo* lymph node imaging under physiological conditions

Using our system, we conducted a biological investigation of a common model system for response to vaccination, *in vivo* imaging of a murine popliteal lymph node (pLN) in anesthetized mouse. Subunit vaccines are a clinically used subset of vaccines in which patients are injected with both a part of a pathogen (the antigen/subunit) and an immunogenic molecule to elicit a protective immune response (the adjuvant). A common model system for these consists of mice being immunized with Ovalbumin (OVA), a protein in egg whites, as a model antigen and lipopolysaccharide (LPS) adjuvant. Before immunization, fluorescently labelled T cells that specifically respond to OVA (monoclonal OT-I and OT-II T cells) are also transferred to the host mouse so that their antigen-specific behavior can be observed in relation to antigen-presenting cells (APCs) in the local lymph node where the initial immune response occurs.

Typically, these experiments can only image a small volume of the lymph node at once. In order to visualize a sufficient number of antigen-specific T cells, previous studies transferred 2-3 orders-of-magnitude more monoclonal cells than would typically exist under physiological conditions, a modification that is well established to alter the dynamics and outcomes of immune responses [5, 6, 7, 8]. With LAMI, we can image 2 orders-of-magnitude larger volumes of tissue (1 vs 10^−2^ mm^3^), so the physiologically unrealistic addition of extra cells is no longer needed. We use an endogenous population of fluorescently labelled APCs, type I conventional dendritic cells (cDC1s) labeled with Venus under the XCR1 promoter [16].

24 hours after immunization with LPS+OVA, the cDC1 network exhibited a marked reorganization (**Fig. S8, Movie S5, S6**), with XCR1+ cells clustering closer to each other and moving from a more even distribution throughout various areas of the lymph node into primarily the paracortex. We found that these clusters of DCs were located primarily around OT-I (CD8 T cells specific to Ova) rather than OT-II (CD4 T cells specific to Ova) or polyclonal T cells, and closer to high endothelial venules than in the control condition.

Imaging and tracking DCs in a control condition and at 24 hours after immunization revealed that this reorganization was accompanied by a change in motility, with DCs at the 24 hour time point moving both more slowly and in a subdiffusive manner, thus confining themselves to smaller areas (**Movie S7**) compared to the more exploratory behavior of the control condition (Fig. 3a). This decrease in average motility appeared global with respect to anatomical subregions and the local density of other dendritic cells (**Fig. S9**). These changes in DC motility were also accompanied by changes in T cell motility in an antigen-specific manner. OT-I T cells, which appeared at the center of dense clusters of DCs, showed the most confined motility compared to polyclonal controls, while OT-II cells were often found on the edges of these clusters with slightly higher motility (Fig. 3b, **Movie S7**).

**Figure 3:**
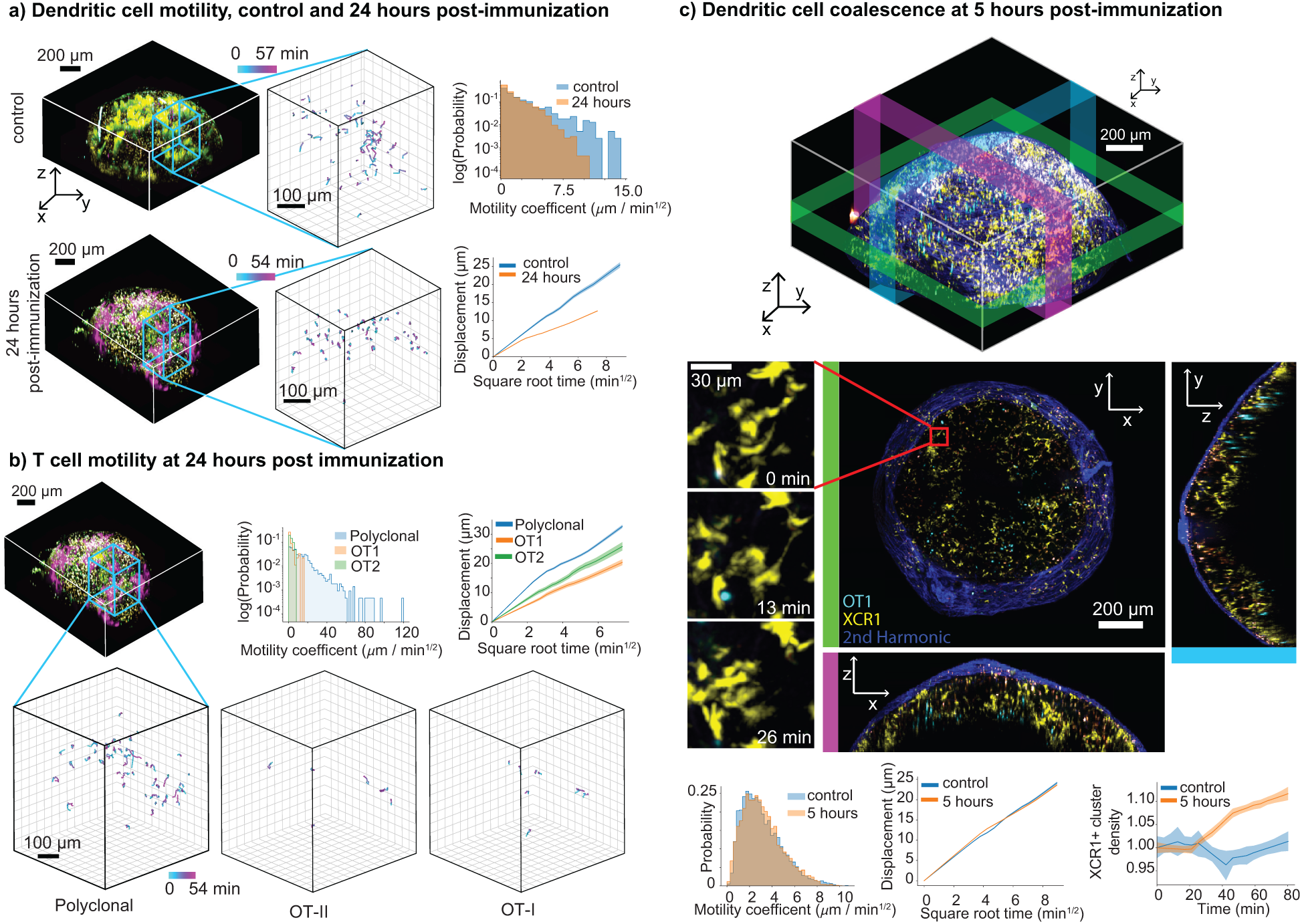
Immune responses under physiological conditions. a) Distinct changes in global behavior of antigen presenting cells as measured by XCR1+ dendritic cell (DC) motility 24 hours after immunization show the cell behavioral correlates of developing immune responses. Tracks of motility in control and 24-hour post immunization (left), log histograms of motility coefficients (right top), and displacement vs square root time (right bottom) show that DCs switch from faster random walk behavior in control (i.e. straight line in bottom right plot) to slower, confined motility 24 hours post immunization. b) T cell motility at 24 hours post-immunization. (Top middle) log histograms of OT1, OT2, polyclonal T cells. (Top right) displacement vs root time plots. (Bottom) tracks of T cell motility. c) DC clustering can be visualized and quantified on whole lymph node level. Top, 3D view with colored bars marking areas shown in 2D projections. b) XY, YZ, XZ projections with zoomed-in area showing example of DC cluster forming over 26 minutes. (Bottom) histograms of DC motility at 5 hours post-infection vs control, displacement vs square root time, averaged normalized density over time in 5 hours post-infection vs control dataset. Error bars on all plots show 95% confidence intervals.

To understand how this reorganization takes place, we next imaged lymph nodes 5 hours after immunization. Although DC motility has not yet changed at this time point, the increasing formation of clusters is detectable on the timescale of an hour (Fig. 3c). Over time, new clusters appeared to form both from spatially separated DCs moving towards one another, and from isolated DCs moving towards and joining larger existing clusters (**Movie S8, S9**).

They also reveal that there is a marked difference in the location and behavior of DC networks encountered by T cells that enter an inflamed lymph node at the beginning versus the later stages of an immune response. Notably, they also show that the larger-scale DC reorganization precedes the T cell activation-induced motility arrest that we and others observe amongst antigen-specific T cells at the 24 hour time point.

We speculate that this increased local concentration of dendritic cells may be necessary for rare, antigen-specific T cells to find one another and form the homotypic clusters required for robust immunological memory [17]. The reorganized environment could be an important factor in the difference in differentiation fate of T cells that enter lymph nodes early vs late in immune responses [18].

## Discussion

Multiphoton microscopy (MPM) is the modality of choice for imaging deep into scattering tissue. Here, we have demonstrated how a computational imaging approach, learned adaptive multiphoton illumination (LAMI), can be used for *in vivo* imaging of lymph nodes with volumes of tissue two orders-of-magnitude larger than existing approaches. This technology opens up new areas of investigation by allowing for more physiological conditions, such as in our case of lower cell frequencies in lymph node imaging.

LAMI is particularly useful for highly-scattering samples with non-flat surfaces (e.g. large organoids or embryos), which have more complicated functions mapping shape to excitation. Applying LAMI to other tissues will require development of sample-specific standard candles. There are many possibilities for these–the only strict conditions are having a labelling density that is not location specific and that individual standard candles can be spatially resolved. Some possibilities include genetically-encoded cytoplasmic fluorophores or organelles or fluorescent beads. Other samples will also require a means of building a map of the sample surface. Though this work uses second harmonic generation signal at the sample surface, reflected visible light might be better suited for this purpose. This process could also be automated to improve imaging speed.

Although scattering of excitation light is likely the largest factor responsible for the drop in fluorescence with depth, absorption also plays a role, especially when imaging deeper into the sample. The fact that far-red fluorphores can be seen at greater depths than those in the visible spectrum supports this fact (e.g. eFluor670 cells in Fig. 2b). The neural network used in LAMI is learning to compensate not only for the scattering of excitation light, but also the absorption of emission light. This implies that greater fluorescence emission, and also greater photobleaching, occurs at greater depths (which we also observed). It is also possible that the sample does not have a spatially-uniform scattering potential, but that the neural network learns to implicitly predict and compensate for this effect.

There are many areas in which LAMI could be substantially improved. The biggest issue in delivering the correct amount of power to each point in intravital imaging of lymph nodes was the map of the sample surface becoming outdated as the sample drifted over time. To combat this, we employed both a drift correction algorithm and periodically recreated the surface in between time points based on the most recent imaging data (**Supplementary materials and methods**). We note that our system used a modified multiphoton system not explicitly designed for this purpose, and building a system from scratch with better hardware synchronization between scanning mirrors, focus, and excitation power would increase temporal resolution several fold and lessen the impact of temporal drift. Using state-of-the art image denoising methods [19] would also allow for faster scanning.

The maximum depth of LAMI in our experiments was limited by the maximum excitation laser power that could be delivered. A more powerful excitation laser could push this limit deeper, or using 3-photon, rather than 2-photon excitation. Another improvement to depth could be made by coupling adaptive illumination with adaptive optics (AO). Incorporating AO could lessen the loss in resolution with depth and potentially restore diffraction-limited resolution deep in the sample. Combining ideas form LAMI with adaptive optics could be especially powerful. One limitation of adaptive optics in deep tissue MPM is the need for feedback from fluorescent sources to pre-compensate for scattering [20, 21, 9], making the achieved correction dependent on the brightness and distribution of the fluorescent source being imaged. We have demonstrated in this work that it is possible to predict the appropriate excitation amplitude from sample shape alone. We speculate that a similar correction might be possible for the phase of excitation light, since scattering is caused by inhomogeneities in refractive index, and the largest change in refractive index seen by the excitation wavefront is likely to be at the surface of the sample when it passes from water into tissue. Deterministic corrections based on the shape of the sample surface have indeed shown to improve resolution in cleared tissue [22], and the additional flexibility of a neural network could provide room for further improvement.

In contrast to contemporary techniques based on deep learning [14], the neural network we employ is simple and shallow (1 hidden layer with 200 hidden units). Adding layers did not increase the performance of this network on a test set. We believe this is a consequence of the relatively small training set sizes we used (10^4^ − 10^5^ examples). Larger and more diverse training sets and larger networks would likely improve performance and potentially allow for additional output predictions such as wavefront corrections.

In conclusion, LAMI is a powerful technique for adaptive illumination in multiphoton imaging, with the potential for opening a range of new types of biological investigations. We were able to implement it on an existing two-photon microscope using only an Arduino-like programmable micro-controller and a small op-amp circuit for less than $50. Furthermore, the software we used is fully open source and compatible with a wide range of microscopes [23].

## Acknowledgements

This project was funded by a Packard Fellowship award to Laura Waller; STROBE: A NSF Science & Technology Center; A Google Cloud research grant award to Henry Pinkard/Laura Waller; an NSF Graduate Research Fellowship awarded to Henry Pinkard; and a Berkeley Institute for Data Science/UCSF Bakar Computational Health Sciences Institute Fellowship awarded to Henry Pinkard with support from the Koret Foundation, the Gordon and Betty Moore Foundation, and the Alfred P. Sloan Foundation to the University of California, Berkeley. The authors thank BIDS and its personnel for providing physical space and general logistical and technical support, Jonathan Shewchuk for helpful conversations during development, and Sandra Baker and Bob Pinkard for support over the course of the project.

## Notes

https://github.com/henrypinkard/LymphoSight

### Competing Interest Statement

The authors have declared no competing interest.

