## Supplementary materials for "Learned adaptive multiphoton illumination microscopy"

**This PDF includes:**

Materials and Methods

Figs. S1 to S9

Captions for Movies S1 to S9

**List of supplementary figures:**


S1: Nonuniform excitation across field on curved tissues

S2: Spatial light modulator test patterns

S3: Spherical tissue ray-optics scattering model

S4: Comparison of excitation strategies

S5: Registration as optimization

S6: Motion artifact correction and active learning-based cell detection

S7: Engineered features for cell classification

S8: Reorganization of lymph nodes 24 hours after immunization

S9: Dendritic cell motility changes in different anatomical locations


**List of supplementary movies:**


S1: Standard candle training data

S2: ex vivo lymph node imaged with adaptive excitation

S3: Correction of transverse movements within Z-stack

S4: 3D volume timelapse before and after motion correction and registration

S5: Cleared lymph node (no immunization)

S6: Cleared lymph node 24 hours after immunization

S7: In vivo immune response 24 hours after immunization

S8: In vivo whole lymph node dynamics 5 hours after immunization

S9: In vivo cell dynamics 5 hours after immunization

### Materials and Methods

Microscope

All imaging was performed on a custom-built two-photon microscope (with 20× 1.05 NA water immersion objective) equipped with two Ti:sapphire lasers, one MaiTai (Spectra-Physics) and one Chameleon (Coherent). The former was tuned to 810nm and the latter was tuned to 890 nm in order to provide a good combination of incident power and excitation for the set of fluorophores used. The microscope had 6 photomultiplier tube detectors in different bands throughout the visible spectrum, giving 6-channel images. All data was collected using Micro-Magellan [7] software to control the Prior Proscan II XY stage and two Z drives, a ND72Z2LAQ PIFOC Objective Scanning System with a 2000 $\mu$m range, which was used to translate the focus during data collection, and a custom built stepper-motor based Z drive, which was used to re-position the sample due to drift in between successive time points. All Z-stacks were collected with 4 $\mu$m spacing was used between successive planes.

Spatial light modulator

Because the appropriate excitation power varies as a function of X, Y, and Z, we need to modulate laser intensity over all of three dimensions. However, typical two-photon microscopes are equipped to only modulate intensity over Z–by changing the laser intensity between different focal planes. Thus, a custom spatial light modulator (SLM) was built to provide the ability to pattern illumination across a single XY focal plane. By applying different 2D patterns at each focal plane, the laser intensity could be modulated across X, Y, and Z. This SLM takes advantage of the scanning nature of multiphoton microscopy (MPM)–that is, the final image is built up pixel-by-pixel in a raster scanning pattern. This scanning pattern is physically created inside of the microscope by the changing the angle of deflection of two scanning mirrors. One of these mirrors operates in resonant scanning mode, oscillating back and forth with sinusoidal dynamics to control X position within the image. The second mirror is a galvanometer which operates with linear dynamics to control the Y position within the image. Both mirrors are controlled by a custom built controller box (Sutter Instruments), which outputs TTL signals corresponding to completion of a single line and completion of a full frame (line-sync and frame-sync, respectively).

The basic operation of the SLM is to take these TTL signals as input, determine where in the field-of-view (FoV) is currently being scanned, and apply appropriate modulation to the excitation laser based on a pre-loaded pattern. The SLM is built from a Teensy 3.2 (a programmable micro-controller) using the Arduino IDE. It connects to the controlling computer via USB, through which a low-resolution (8x8) XY modulation pattern is pre-loaded via serial communication. The SLM is also connected to the mirror controller's frame-sync and line-sync TTL signals. Each time one of these signals is received, an interrupt fires, which initiates a corresponding timer. In between interrupts, current scanning position in X and Y is determined based on the elapsed time on these timers (using the appropriate inverse cosine mapping for the resonant scanner). The laser modulation is then determined for that point by bilinear interpolation of the low-resolution pattern. This ensures the ability to apply a smooth gradient of excitation across the field rather than a discretized one determined by the resolution of the supplied pattern.

The excitation laser's amplitude is controlled by an electro-optical modulator (EOM), which takes a logic-level input of 0-1.2V (where 0V is off and 1.2V is full power). The modulator's input is controlled by the Teensy's onboard digital-to-analog converter (DAC) via a voltage divider and voltage buffer. The DAC output is put through a voltage divider to lower the logic level from 3.3V to 1.2V in order to utilize the full 12-bit analog control, and the signal is then run through a LM6142 rail-to-rail operational amplifier in a buffer configuration to isolate the DAC output from any downstream current-draw effects.

To validate the performance of the SLM, a uniform fluorescent plastic slide was imaged with different patterns projected onto it (**Fig. 2**). Figure 2a shows a checkerboard pattern, which is not a realistic pattern that would be projected into a lymph node, but demonstrates the resolution capabilities of the SLM. Along the vertical axis, the pattern can be precisely specified on a pixel-by-pixel basis. However, the pattern is blurred along the horizontal direction, resulting from the average of many images, each with a noisy pattern along that dimension due to the resonant scanning mirror moving along the horizontal axis faster than the vertical axis. The fundamental limitation is the clock speed of the Teensy, which limits how fast the voltage to the EOM that modulates the excitation laser can be updated. However, in practice, this noise is not a problem because the excitation power needed is a smoothly varying function, and thus a more realistic pattern for imaging into a sample is a gradient pattern (Fig. 2c).

Imaging experiment setup

The popliteal lymph node was surgically exposed in an anesthetized mouse. Because of the geometry of our surgical setup, only one half of the popliteal lymph node was visible (i.e. the axis running from top of cortex to medulla was perpendicular to the optical axis). Although we were able to image this half of the lymph node, to get a better view of the whole cortical side of the lymph node, we had to cut the afferent lymphatic, so that the lymph node could be reoriented with it's cortex facing the objective lens. The efferent lymphatic and blood vessels were left intact. We note that a better surgical technique might be able to circumvent this limitation.

To start the experiment, the microscope was focused to a point on the top of the lymph node cortex using minimal excitation power and the signal visible from second harmonic generation (SHG). Micro-Magellan's explore mode was then used to rapidly map the cortex of the lymph node using a low excitation power, and interpolation points were marked on collagen signal from the SHG image. This surface was used not only to predict the modulated excitation power, but also to guide data acquisition: Using Micro-magellan's distance from surface 3D acquisition mode, only data within the strip of volume ranging from 10 $\mu$m above the lymph node cortex to 300$\mu$m was acquired, rather than the cuboidal volume bounding this volume. This avoided wasting time imaging areas that were either not part of the lymph node, or so deep within it that they are below the depth limit of 2-photon microscopy. Over time the volume being imaged tended to drift. This was partially compensated for by using the drift correction algorithm described below. However, we limited the use of this algorithm to drift in the Z direction where drift tended to be the most extreme (presumably because of thermal effects or the swelling of the lymph node itself). For XY drift, or for Z drift the algorithm did not correct, we periodically paused acquisition and marked new interpolation points on the cortex of the lymph node, in order to update both the physical area being imaged, and the automated control of the excitation laser.

Image denoising

All data were denoised using spatio-temporal rank filtering [7]. Two full scans of each field-of-view were collected at each focal plane before being fed into a 3×3 spatial extent rank filter. Because of the computational load of performing all the sorting operations associated with this filtering at runtime, a computer with a AMD RYZEN 7 1800X 8-Core 3.6 GHz processor was used for data collection, and the filtering operations were paralellized over all cores. In addition, the final reverse-rank filtering step was done offline to save CPU cycles during acquisition. Before processing data, an additional filtering step using a 2D Gaussian with a 2-pixel sigma kernel was applied to each 2D slice to improve signal-to-noise on downstream tasks. We note that while spatio-temporal rank filtering was designed specifically for the task of cell detection applied here, there may be room for further improvement of real-time denoising (and thus lower doses of excitation light) strategies based on deep learning [11].

Ray optics spherical excitation model

On the way to developing standard candle calibration, we experimented with a simulation framework in which lymph nodes were modeled as spheres with homogeneous scattering potential. This framework had several disadvantages, namely that it was too computationally intensive to run in real time, relied on parameters which had to be heuristically tuned, didn't consider the attenuation of fluorescence emission as it travels back to the detector, and was fundamentally mismatched from actual lymph nodes, which are curved but often far from spherical. However, it was useful as a starting point for calculating the random excitation powers that were applied to generate the standard candle training data (even though this may not have been absolutely necessary for generating random excitation). It is also useful to understand why it is difficult to accurately make a physics-based model of this problem, and why machine learning is especially useful. The details of this model are described below.

For a single ray propagating through tissue the proportion of photons which remain unscattered and the two-photon fluorescence intensity decay exponentially with depth[2]: $F = P_0 e^{-\frac{2z}{l_s}}$, where $F$ is the two-photon fluorescence excitation, $z$ is the distance of propagation, $P_0$ is the incident power, $l_s$ is the "mean free path" for a given tissue at a given wavelength, which measures the average distance between scattering events.

We assume a beam with a Gaussian profile at the back focal plane of an objective lens, which implies that the amplitude and intensity of the cross-sectional profile of the focusing beam are also both Gaussian. We also assume the contribution from photons that are multiply scattered back to the focal point is negligible and that scattered light does not contribute to the two-photon excitation at the focal point. Using a geometric optics model with these assumptions, the attenuation of each ray propagating towards the focal point can be considered separately. Thus, we can calculate the amount of fluorescence emission at the focal point by numerically integrating over all rays in the numerical aperture of the objective, with a known tissue geometry and scattering mean free path (**Fig. 3**a). Figure 3b shows the output of such a simulation for a spherical lymph node of a given size. Relative excitation power is that factor by which input power would need to be increased to yield the same fluorescence as if there were no scattering. It is parameterized by the vertical distance from the focal point to the lymph node surface, and the normal angle at that surface.

Standard candle calibration

The training data for standard calibration was collected by imaging an inguinal lymph node *ex vivo*, which had previously been seeded with $2 \times 10^6$ lymphocytes from a Ubiquitin-GFP mouse and 2x$10^6$ lymphocytes from a B6, which had been labelled in vitro with eFluor-670 (e670). Each population was used as a standard candle for one of the two excitation lasers on the system, which had their wavelengths tuned to 810nm and 890nm. Two separate images were recorded, one with each laser on. The lymph node was imaged by tiling multiple Z-stacks in XY to cover the full 3D volume. Each Z-stack was imaged with power determined using the output of the spherical model described above, multiplied by a randomizing factor drawn from a uniform distribution between 0.5 and 2. This randomly distributed brightness data was then fed into the cell segmentation and identification pipeline described below. The mean brightness was taken for all voxels within each segmented region as the brightness of the standard candle. The standard candle's spatial location was used to determine the EOM voltage applied at that point in space, its location in the XY FoV, and a set of statistics to serve as effective descriptors of the physics of light scattering and emission light absorption.

The physical parameters were computed by measuring 12 distances from the focal point of the standard candle to the top of the interpolation marking the cortex of the lymph node (**Fig 1** in main text). All 12 distances were measured along directions that had the same angle of inclination to the optical axis ($\phi$), with equally spaced rotations about the optical axis. Taken in its raw form, each element of this feature vector is associated with a specific absolute direction in the coordinate space of the microscope. The microscope should be approximately rotationally-symmetric about its optical axis. We don't want the machine learning model to have to learn this symmetry from data, because it would needlessly increase the amount of training data needed. Thus, we explicitly build in this assumption by binning all distances into a histogram. We use nonlinearly-spaced bin edges for this histogram, based on the intuition that scattering follows exponential dynamics with propagation distance, so relative difference in short distances of propagation are more significant than those same differences at long distances. The bin edges of this histogram were calculated by taking 13 equally-spaced points from zero to one, and putting them through the transformation $f(x) = (x^{1.5})(350\mu m)$, where 350 $\mu$m is the propagation distance beyond which we don't expect excitation light to yield any fluorescence excitation.

Standard candle brightness, location in XY FoV, and the physical parameter vector were concatenated into a single feature vector. Each of these feature vectors corresponded to one standard candle cell and was associated with a scalar that stored the voltage of the EOM used to image that standard candle. The total number of these pairs were 4000 for the GFP standard candles and 14000 for the e670 standard candles. We standardized all feature vectors by subtracting their element-wise mean and dividing by their element-wise standard deviation. We then trained a fully connected neural network with one 200-unit hidden layer and a single scalar output. The network was trained using the Adam optimizer, dropout with probability 0.5 at training, and a batch size of 1000. Training was continued until the loss on the validation set ceased to decrease.

The output of this network is the voltage on a particular EOM. Because the goal of this network is to deliver the right amount of excitation power, as opposed to voltage, we converted this voltage into an estimate of relative excitation power (in arbitrary units) before feeding it into a squared error loss function. We measured the function relating EOM voltage to incident power empirically by placing a laser power meter at the focal plane of the objective lens and measuring the incident power under several different voltages. We found this curve to be well approximated by a sinusoid, so we fit the parameters of this sinusoid and used it directly in the loss function.

We experimented with several different architectures before finding the one that worked best with our data. Neither adding additional hidden layers, nor increasing the width of the existing hidden layer beyond 200 improved performance. The best performing value of $\phi$ (The angle of inclination to the optical axis) was 20°. Neither other angles, nor using multiple angles improved performance on the validation set. This was somewhat surprising, as we would have expected more information about the local geometry to improve prediction. We suspect that this might be the case with a larger training set.

#### Using standard candle calibration to control laser power

On later experiments, we loaded the trained weights of the network, computed its output for 64 points in an 8×8 grid for each XY image, and sent these values to the SLM through serial communication. The element of the vector corresponding to standard candle brightness must be chosen manually, and can be thought of a z-score of the distribution of brightnesses in the training set (since the training set was standardized prior to training). For example, picking a value of 0 means the network will provide the right laser power to achieve the mean brightness in the training set. Picking a value of -1 means it will aim for a brightness 1 standard deviation below the mean value of the training set.

To calculate the physical parameter feature vector, we computed the interpolation of the lymph node surface as described previously. This interpolation yields a function of the form $z(x, y)$, where there is a single z coordinate for every XY position (unless the XY position is outside the convex hull of the XY coordinates of all points, in which case it is undefined). To avoid having to repeatedly recalculate this function, it is evaluated automatically over a grid of XY test points and cached in RAM by Micro-Magellan. In order to fill out the physical parameter feature vector, we must calculate the distance from an XYZ location inside the lymph node to it's intersection with the interpolated surface. We measure this distance numerically, using a binary search algorithm. This algorithm starts with a value larger than any distance we expect to measure (i.e. 2000$\mu$m), tests whether the Z value for this XY position is above the surface interpolation or undefined (which means it is outside the lymph node), halves the search space, and repeats this test until the distance is within some tolerance (we used $\mu$m). These calculations were all handled on a separate thread from acquisition so that they could be pre-computed and not slow down acquisition. We note that our strategy of sending each pattern out as a serial command certainly prevents the system from running as fast as it might otherwise be able. Sending out many such patterns at once and relying on a system that uses hardware TTL triggering should dramatically increase the temporal resolution of this technique.

Validating standard candle calibrated excitation

To validate the use of standard candle calibrated excitation, we transferred $2\times10^6$ GFP lymphocytes, $2\times10^6$ RFP lymphocytes, and $2\times10^6$ e670 lymphocytes and imaged its mediastinal lymph node *ex vivo*. We note that the mediastinal lymph node is quite different in size and shape than an inguinal lymph node. The lymph node was imaged with constant excitation, excitation predicted by the spherical ray optics model, and excitation predicted by the standard candle neural network (**Fig. 4**). The transferred lymphocytes included both T cells and B cells, meaning that there should be fluorescently labelled cells throughout the volume of the lymph node. A detailed comparison of the resultant images is shown in Fig. 4, which also shows that the learned excitation outperforms the spherical-ray optics model.

Drift correction

Focus drift, primarily in the Z direction, was present in all experiments at rates on the order of $1\mu$m/min. This is unsurprising given the massive influx of cells to lymph nodes during inflammatory reactions. It was essential to compensate for this drift, because not doing so would lead to a mismatch between the coordinates of our interpolation marking the lymph node cortex and it's actual location, which would in turn mean the automated excitation would be misapplied. To compensate for this drift, we designed a drift compensation algorithm that ran after each time point, and changed the Z position of secondary Z focus drive (i.e. not one used to step through Z-stacks) after each time point. Estimates of drift were based on the second harmonic generation signal from the fibers in the lymph node cortex, which were a convenient choice because their contrast was not dependent on fluorescent labelling, and their spectral channel (violet) had relatively little cross-talk with other fluorophores. At each time point after the second, the cross-correlation of the 3D image in violet channel was taken with the corresponding image from the previous time point. The maximum of this function was taken within every 2D image corresponding to a single slice, and a cubic spline was fit to these maxima. The argmax of the resulting smooth curve was used to estimate the offset in Z between two successive timepoints with subpixel accuracy. This estimate was used to update an exponential moving average that estimated the rate of drift, so that both the existing drift from the previous timepoint could be corrected, and the expected future drift could be pre-compensated for. In practice, this algorithm worked well enough to stabilize the sample enough for the adaptive illumination to be correctly applied. Remaining drift in the imaging data was corrected computationally as described below.

Image registration

We identified three types of movement artifacts that occurred during intravital imaging. 1) Due to the mouse's breathing, there were periodic movements of successive images relative to one another within each Z-stack. These movements could be well approximated by motion within the XY plane, in part because of the geometry of the imaging setup, and in part because of the heavily anisotropic resolution of the imaging system, in which objects were blurred out along the Z axis much more so than X and Y. 2) Individual Z-stacks were misaligned with each other in X, Y and Z. This seemed likely to be caused by physical movement of the sample as a result of some combination of thermally-induced focus drift and biological changes leading to small tissue movements. 3) Global movements of the entire sample over time. All three remained to some degree even after experimental optimizations to improve the system stability and pre-heating the objective lens to minimize thermal drift.

To correct these artifacts, we used a three stage procedure with each step corresponding to a type of movement artifact. Although cross-correlation is often the first choice for rigid image registration problems in the literature, it was found to be ineffective for solving two of the three of problems. Thus, we employed a more general framework, using iterative optimization to compute *maximum a posteriori* (MAP) estimates. This framework depends on the ability to transform and resample the image in a differentiable manner. As shown in Fig. S5a, we can set up a general image registration problem that can be solved by numerical optimization by creating a parametric model for how pixels move relative to one another, resampling the raw image based on the current parameters of this model, and then computing a loss function that describes how well the alignment based on this transformation is. This paradigm enables us to solve general MAP estimation problems of the following form with iterative optimization:

$$\boldsymbol{\theta}^* = \arg\min_{\boldsymbol{\theta}} L(f_1(\boldsymbol{\theta}), f_2(\boldsymbol{\theta}), ...) + R(\boldsymbol{\theta})$$

Where $\boldsymbol{\theta}$ are the parameters to optimize, $f_n$ is the transformation and resampling of the $n^{th}$ subset of pixels, $L$ the loss function, and $R(\boldsymbol{\theta})$ is a regularization term for the parameters that allows incorporation of prior knowledge. We used the deep learning library TensorFlow to set up these optimization problems. This had the advantage of being able to automatically calculate the derivatives needed for optimization using built-in automatic differentiation capabilities. Often, these problems used extremely large amounts of RAM, because all image pixels were stored in memory when performing optimization. We were able to do this by using virtual machines on Google Cloud Platform with extremely large amounts of memory (>1TB). However, we note that it would be possible to reduce the RAM requirements by downsampling the images, or more carefully coding the optimization models to only use relevant parts of the images rather than every pixel.

For the first correction, movements in XY for each Z-stack, each Z stack was optimized separately. We observed that XY movements were almost always confined to a single z plane and that looking at an XZ or XY image of the stack, these movements were clearly visible as discontinuities along the Z axis. Thus we parameterized the model by a (number of Z-planes)×2 vector, corresponding to an XY shift for each plane. For this correction, all channels except for the channel corresponding to second harmonic generation were used. The loss function was taken as the sum over all pixel-wise mean-squared differences between consecutive z planes, normalized by the total squared intensity in the image (which was necessary to ensure that the learning rate of the optimization did not need to be adjusted to accommodate the total brightness of the Z-stack):

$$L(\boldsymbol{x}, \boldsymbol{y}) = \frac{\sum_{j=0}^{N-1} \sum_{x',y'} (\boldsymbol{I}(x' + x_j, y' + y_j, z_j) - \boldsymbol{I}(x' + x_{j+1}, y' + y_{j+1}, z_{j+1}))^2}{\sum_{x',y',z'} \boldsymbol{I}(x', y', z'_j)^2},$$

where $\boldsymbol{x}$ and $\boldsymbol{y}$ are vectors holding the translations at each slice, $j$ is the index of the z plane, $N$ is the total number of Z planes, $\boldsymbol{I(x, y, z)}$ is a pixel in the raw Z-stack, and $x'$, $y'$ are the coordinates of pixels in the raw image. The regularization in this problem was a quadratic penalty on the sizes of the translations (implicitly encoding a prior that these translations should be normally distributed about 0) multiplied by an empirically-determined weighting factor:

$$R(\boldsymbol{x}, \boldsymbol{y}) = \lambda(\|\boldsymbol{x}\|_2^2 + \|\boldsymbol{y}\|_2^2)$$

The value of lambda used was $8{\times}10^{-3}$. The model was optimized using the Adam optimizer and a learning rate of 1. Optimization proceeded until a the total loss had failed to decrease for 10 iterations.

The second correction, fixing movements over time, was computationally much easier to solve, because the strong signal of the similarity between consecutive time points in channels made registration not especially difficult if the correct channels were used. For this reason, this correction did not require iterative optimization, and could instead be solved with cross-correlation alone. The 3D cross-correlation was taken between every consecutive two time points for each Z-stack. The location of the maximum value of each of these cross-correlations gave the optimal 3D translation between consecutive time points, and taking a cumulative sum of these pairwise shifts gave an absolute shift for each stack over time.

The third correction, finding the optimal stitching alignment between each Z-stack, was the most computationally challenging of these problems. This is because it has a relatively small amount of signal (i.e. the overlapping areas of each Z-stack, which was less than 10% of the total volume of each Z-stack). Furthermore the signal in these areas was relatively weak, because it was most susceptible to photo-bleaching since it is exposed to excitation light multiple times at each point. Furthermore, since stacks were often taken a few minutes apart, the content in these overlapping regions often changed. Compensating for these difficulties not only required using the iterative optimization framework with an appropriate loss function that accounted for variations in image brightness and proper regularization, but also carefully choosing which channels to use registration based on the presence of non-motile fiducial signals. Most of the datasets we collected had a channel with high endothelial venules, a large and immobile structure in the lymph node, fluorescently labelled, and these channels were often the most useful due to strong signal and lack of movement. We also found good performance by including the second harmonic generation channel that provided signal from the collagen fibers in the lymph node cortex. Finally, we noticed that autofluorescent cells were numerous and immobile throughout the lymph node. Because autofluorescence has a broad emission spectrum that appears across 3-4 channels at once, as opposed to the labelled structures which appear over only 1-2, we were able to isolate the signal from these cells by taking the minimum pixel value over several channels.

As shown in Fig. 6b, each Z-stack was parameterized by a 3 element vector that corresponded to its X,Y, and Z shifts. The loss function was taken as the mean of all of the correlation coefficients of the pixels in the overlapping regions of every pair of adjacent Z-stacks. Correlation coefficients are a better choice of loss for this task than cross-correlation, because they better account for variations in image brightness [5]. Optimization was performed using Newton's method with a trust region constraint. Rather than performing optimization on each time point separately, all time points were averaged together and a single optimization was performed taking all information into account. This was possible because relative movements between stacks that differed by time point had already been corrected by cross-correlations in step 2. Because of the strong signal afforded by averaging multiple time points together, no regularization was needed.

Cell identification–feature engineering

Cells were detected in a two-stage pipeline that first utilized 3D segmentation to identify cell candidates, followed by machine learning to classify which of those cell candidates belonged to a population of interest. Candidate regions were generated using the segmentation algorithm (Fig. 2c in main text) built into Imaris 7.6.5 (i.e. the "surfaces" module), which includes a filtering step to smooth the data, a local background subtraction step to account for variations in brightness, a thresholding step to generate segmented regions, and a splitting step, in which seed points of a certain size are generated, and segmented regions are split based on these seed points. Candidates were generated for each population of interest (i.e. each fluorescent label) through the ImarisXT Matlab interface. Next, each candidate region was "featurized" by computing a set of descriptive statistics about the pixels enclosed within it. By default, Imaris outputs a set of 97 such statistics for each candidate, including intensity means, standard deviations, minimums, maximums, as well as a number of morphological features. However, these features are specific neither to the biological or technical context of the data, and we found them to not be effective in all cases for training high-quality classifiers. Thus, we engineered a set of additional features to better capture the variations that are useful for classifying cells.

First, we reasoned that since all spectral channels are collected simultaneous in two-photon microscopy, the ratios of intensity in different channels contains important information. Treating each set of spectral statistics (e.g. intensity means for different channels) as a 6-dimensional vector (for a 6-channel image), we subtracted the background pixel value for each channel, and normalized to unit length. This "spectral normalization" takes advantage of the fact that that intensity measurements for a given fluorescent object are all proportional to the excitation power delivered to the focal point, and thus it normalizes intensity statistics while preserving their ratio. It also creates an additional feature from the magnitude of the vector prior to normalization, which captures the brightness of the object irrespective of it spectral characteristics.

We also designed several feature classes based on the observation that one of the failure modes of the segmentation algorithm in the candidate generation step was that it often created a single region around a cell of interest along with a second cell in close contact to it that expressed a different fluorophore, but had spectral bleed-through into the channel on which the segmentation was run. Thus, intensity weighted centers-of-mass (COMs) within each region would be expected to show greater variance among the different spectral channels compared to a surface that surrounds a structure single source of fluorescence intensity. This should hold true even if the the two objects surrounded by a single surface shared emission in the same channels, as long as the spectral profile of the two objects differs. With this in mind, we computed features for all pairwise distances between the intensity-weighted COM for different channels, as well as the distance from each intensity-weighted COM to the non-intensity weighted COM. With the same reasoning, we also added the correlation matrices containing the pairwise correlations between channels for all pixels within each candidate region as features (Figure S5a).

Finally, to more directly address the issue of overlapping, spectrally dissimilar cells (which are often the most biologically interesting case), we designed an algorithm to identify sub-regions of pixels within each candidate region that has a spectrum that is most similar to a reference spectrum (i.e. the spectrum of the fluorophore of the cell of interest). This algorithm is based on the normalized cut segmentation algorithm [9]. However, unlike that algorithm, which is designed for use on grayscale images, and builds an adjacency matrix for all pixels based on a combination of their spatial and intensity differences, our algorithm segments regions based on differences in their spatial and spectral distances. This is accomplished by defining distances between each pair of pixels as:$d_{i,j} = \alpha \|\mathbf{r}_i, \mathbf{r}_j\|_2^2 + \beta \hat{\mathbf{s}}_i^T \hat{\mathbf{s}}_j$, where $\mathbf{r}_i$ and $\mathbf{r}_j$ are the spatial coordinates of the two pixels, $\hat{\mathbf{s}}_i$ and $\hat{\mathbf{s}}_j$ are their unit norm intensity vectors across all channels, and $\alpha$ and $\beta$ are tuning parameters. Then, an adjacency matrix can be constructed by defining the adjacency between pixel $i$ and pixel $j$ as $w_{i,j} = e^{-d_{i,j}}$. The spectral clustering method defined in [6] can then be used to break all pixels into distinct regions, and the normalized cut region of interest (NC-ROI) most similar to a given reference intensity can be used for further downstream processing. Using this method, a number of additional features were calculated for the pixels within each NC-ROI.

To validate that these features were in fact useful, we performed two types of analyses. First, we looked at which types candidates cells the classifier repeatedly failed to correctly classify. By running k-fold cross validation on a ground truth set of labelled candidates, we were able to identify which candidates the classifier failed to correctly classify. Overlaying these results on principal component analysis plot of the spectral variation among the cells of interest (RFP labelled T cells), we found that the misclassified cells were often spectral outliers as a result of spatial overlap with some other fluorescent structure (Fig S6, left). However, including the engineered features in this experiment dramatically reduced the miscalssificaiton of these cells.

Next we ran a bootstrap analysis to identify the most useful features. We performed regularization and variable selection via the elastic net procedure. Elastic net is a useful method for identifying a sparse subset of useful predictors in a dataset with correlated predictors.

For the dataset examined, the number of labeled T cells was significantly lower than the number of non T cells (T cells: 204, non T cells: 38, 575). In order to keep a balanced training and test dataset, we partitioned the data so that the model performs training and testing on similar sample distributions. In particular, we performed 100 bootstrap resampling procedures from both T cell and non T cell data of approximately equal sample size. The model obtained was further tested on a smaller test dataset with equal T cell and non T cell ratio, set aside at the beginning of the procedure.

For each bootstrapped sample set, we further ran 1000 iterations of randomly picking test set/training set partitions. This step was performed in order to asses the stability and overall distribution of the lambda parameter picked for each model. Despite lambda being chosen through 5-fold cross-validation procedure, it is still specific to the prior decided training/test set partition. By performing additional random partitions of the bootstrapped dataset, we can break this dependence. Additionally, we can also look at the distribution of cross-validation error provided by the glmnet package, as well as the misclassification error from the test set. The final model was fitted using the lambda parameter with the lowest cross-validation and misclassification error for the bootstrapped sample set. We looked at the averaged probabilities across all the samples, as well the total number of times each predictor was chosen by the elastic net out of 100 bootstrap subsamples.

The final results of this analysis can be seen in Figure S6d. Many of the engineered features are among the strongest predictors, further validating their usefulness for this task.

Cell identification – active learning

Having developed a useful set of task-specific engineered features that can train high quality classifiers with sufficient labelled training data, the last remaining piece of the pipeline is a scheme for generating labelled training data. Often, this can be the most time consuming piece of developing a machine learning system. To alleviate this bottleneck, we drew from the field of active learning, a paradigm in which a classifier chooses which data points receive labels, allowing them to learn more efficiently by seeing more informative examples [3] . Specifically, we use the strategy of ”uncertainty sampling” [4], in which the classifier outputs a number between 0 and 1 for each example (with 0 being complete certainty of one class and 1 being complete certainty of the other), and the example with a value closest to 0.5 is then selected and sent to a human for labelling. This process is then repeated until enough training data is labelled to train a classifier that generalizes well to the remaining unlabelled data.

Applied specifically to the problem of classifying which candidate regions corresponded to cells, the workflow was a follows: After computing the engineered features for all candidate regions, we first labelled one example of a candidate that belong to the population and one that did not. These labels were used to train the classifier (a small, fully connected neural network with 12 hidden units). Because classification accuracy usually increased by averaging the predictions of multiple neural nets, 3 were trained in parallel and their predictions were averaged. When making final predictions of cell populations, 100 neural nets were averaged. The neural net was trained in Matlab, and the labelling interface for selecting cells was built with Imaris for data visualization, and Matlab script on the backend that communicated interactively with Imaris through the ImarisXT interface. We periodically predicted the identities of all candidates, in order to identify particularly difficult examples even faster and manually label them.

Although well justified from a theoretical standpoint as a means to make exponential gains in data labelling efficiency on idealized problems [8], there remained the question of whether active learning had the same effect on this problem. To answer this, we generated a ground truth set of labels by carefully manually labelling every cell on a limited dataset of candidates for RFP-labelled T cells. We reviewed each cell multiple times to be sure that its label was not a false positive, and searched manually through the data volume to identify any false negatives. Next, one positive and one negative candidate were randomly selected and assigned labels. This labelled set was used to simulate the uncertainty sampling procedure, drawing labels from the ground truth set rather than a human labeller. The accuracy on the remaining unlabelled examples was used to assess performance. As the plot in Figure S6d demonstrates, uncertainty sampling vastly outperformed random sampling for this classification task.

Cell tracking and statistical analysis

All data analysis was performed in using custom scripts written using tools from the Scientific Python stack [10]. Curves were fit to the means of scatterplot data (e.g. displacement vs. square root time plots) using locally weighted scatterplot smoothing (LOWESS) using locally linear regression [1] using a tricubic or Gaussian weighting functions. Sigma and alpha parameters were tuned manually for each comparison to capture the data while smoothing out noise. Error bars represent 95% confidence intervals derived from bootstrap resampling of data with 500 iterations

XCR1+ cluster density was computed by counting the number of other XCR1 cells within 100 $\mu$m of each detected XCR1+ cell divided by the total number of XCR1 cells detected at each time point. Cells were tracked over multiple time points using the Brownian motion tracking algorithm in Imaris 7.6.5.

Mouse Strains

All mice were treated in accordance with the regulatory standards of the National Institutes of Health and American Association of Laboratory Animal Care and were approved by the UCSF Institution of Animal Care and Use Committee (IACUC approval: AN170208). All mice were purchased for acute use or maintained under specific pathogen-free conditions at the University of California, San Francisco Animal Barrier Facility. Mice of either sex ranging in age from 6 – 12 weeks were used for experimentation.

The following mice were purchased from the Jackson Laboratory or bred to a C57BL/6 background: TCRa knock-out, LCMV P14 specific transgenic mice (MMRRC stock no:37394-JAX), C57Bl/6J (stock no:000664), OT-1 (stock no:003831) OT-2 (stock no:004194) transgenic mice interbred with CD2-RFP or ubiquitin-GFP (stock no:004353), and XCR1-Venus mice.

Mouse Immune challenge

OTI and OT2 cells were isolated from lymph nodes of mice. Additionally, polyclonal (C57BL/6) or LCMV P14-specific CD8+ T-cells were isolated as negative controls. Selection was carried out with a negative selection EasySep mouse CD8+ or CD4+ isolation kit (STEMCELL Technologies, 19853 and 19852). If T-cells did not have a transgenic reporter (CD2-RFP or ubiquitin-GFP), they were fluorescently labelled with one of eFluor 670 (ThermoFisher Scientific, 65-0840-85), Violet Proliferation Dye 450 (BD, 562158), or CMTMR (Thermo Fisher Scientific, C2927). Dyes were diluted 1000-fold and incubated with isolated cells for 10-15 minutes minutes in a 37C, 5 percent CO2 incubator. T cells were injected retro-orbitally (r.o.) into Xcr1-Venus recipient mice in 50-100 uL volumes. The number of OT1 and OT2 transferred was 5e4 except for experiments conducted at 5 hours post-infection, where 5e5 cells were transferred in order to visualize more T cell-dendritic cell interactions; for each experiment, equal numbers of OT1 and OT2 cells were transferred. 1e6 control T-cells were transferred. Mice were given a 30 uL footpad injection containing 2.25 ug LPS (Sigma-Aldrich, L6529-1MG) and 20 ug OVA protein (Sigma, A5503-1G) 1-4 days after T-cell transfer; a 30 uL footpad injection of DPBS was used as a negative control to the infection model. To visualize high endothelial venules, in some imaging experiments, 15 ug Meca-79 Alexa Fluor 647 (Novus Biologicals, NB100-77673AF647) or Alexa Fluor 488 (Novus Biologicals, NB100-77673AF488) was transferred r.o. in a volume of 50 uL immediately before imaging.

**Supplementary movie 1: Standard candle training data** Inguinal lymph node seeded with GFP lymphocytes imaged with randomized excitation power to generate standard candle training data

**Supplementary movie 2: ex vivo lymph node imaged with adaptive excitation** Inflamed mediastinal lymph node with adoptively transferred lymphocytes labelled with GFP, RFP, eFluor670 imaged using adaptive excitation.

**Supplementary movie 3: Correction of transverse movements within Z-stack** single Z-stack with tranverse movement artifact caused by breathing (left) and with optimized corrections (right)

**Supplementary movie 4: 3D volume timelapse before and after motion correction and registration** Timelapse before and after registration, motion corrections and stitching

**Supplementary movie 5: Cleared lymph node (no immunization)** Cleared lymph node showing positioning of XCR1+ cells (yellow) when in uninfected condition

**Supplementary movie 6: Cleared lymph node 24 hours after immunization** Cleared lymph node showing positioning of XCR1+ cells (yellow) 24 hours after immunization with LPS

**Supplementary movie 7: In vivo immune response 24 hours after immunization**

**Supplementary movie 8: In vivo whole lymph node dynamics 5 hours after immunization**

**Supplementary movie 9: In vivo cell dynamics 5 hours after immunization and tracked dendritic cells showing clustering behavior**

[10] Pauli Virtanen, Ralf Gommers, Travis E Oliphant, Matt Haberland, Tyler Reddy, Stéfan J Van Der Walt, Matthew Brett, Joshua Wilson, and K Jarrod Millman. computing in Python. *Nature methods*, 2020.

[11] Martin Weigert, Uwe Schmidt, Tobias Boothe, Andreas Müller, Alexandr Dibrov, Akanksha Jain, Benjamin Wilhelm, Deborah Schmidt, Coleman Broaddus, Siân Culley, Mauricio Rocha-Martins, Fabián Segovia-Miranda, Caren Norden, Ricardo Henriques, Marino Zerial, Michele Solimena, Jochen Rink, Pavel Tomancak, Loic Royer, Florian Jug, and Eugene W. Myers. Content-aware image restoration: pushing the limits of fluorescence microscopy. *Nature Methods*, 15(12):1090–1097, 2018.

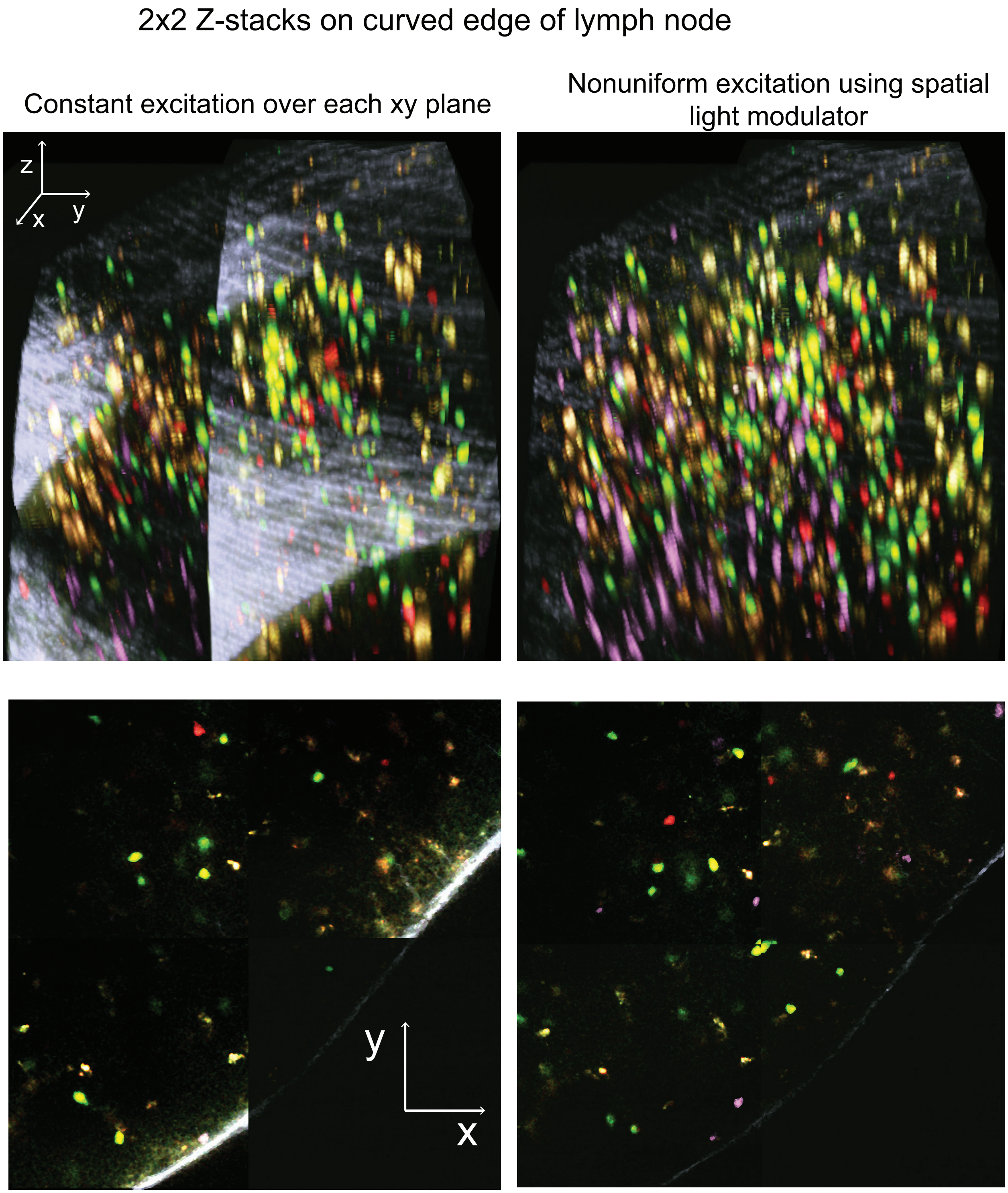


**Figure 1: Nonuniform excitation across field on curved tissue**. Imaging into to curved tissue such as the edge of a lymph node requires variable excitation over the XY field of view. 3D view (top) and 2D slice (bottom) of a 2x2 grid of Z-stacks. Left, constant excitation power within each XY plane in each Z stack. Right, variable excitation power allows excitation to be set correctly for each point in XYZ field of view.

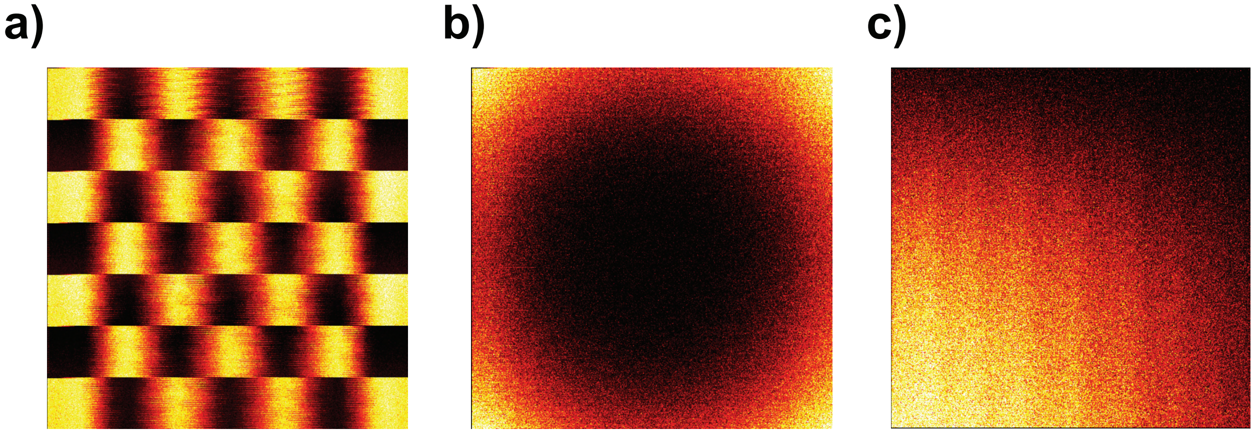


**Figure 2: Spatial Light Modulator Test Patterns.** Images taken on flat fluorescent test slide with different patterns of excitation light. a) A checkerboard pattern demonstrating the difference in horizontal vs. vertical resolution. b) A vignetting compensation pattern, with more excitation at the edges of the field of view. c) A gradient across the field of view pattern.

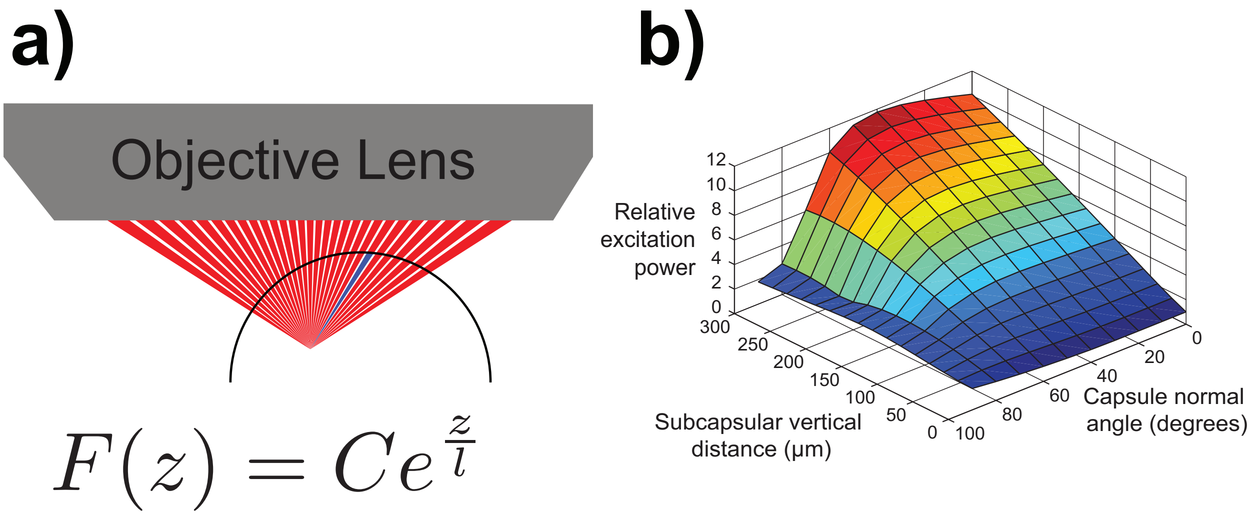


**Figure 3: Spherical tissue ray-optics scattering model.** A previous scattering model used on the way to developing standard candle calibration. In this model, the tissue is assumed to be a sphere with homogeneous scattering potential. a) Fluorescence at the focal point is computed by integrating the contribution from every ray within the cone of the objective's numerical aperture. The contribution of each ray drops off with its propagation distance through tissue ($z$) as shown in the equation. b) The predictions of the model with parameters estimated for lymph node tissue. Relative excitation power is the inverse of the fraction of input power that makes it to the focal point. It is indexed by the vertical distance from the focal point to the top of the tissue, and the normal angle of the sphere directly above the focal point.

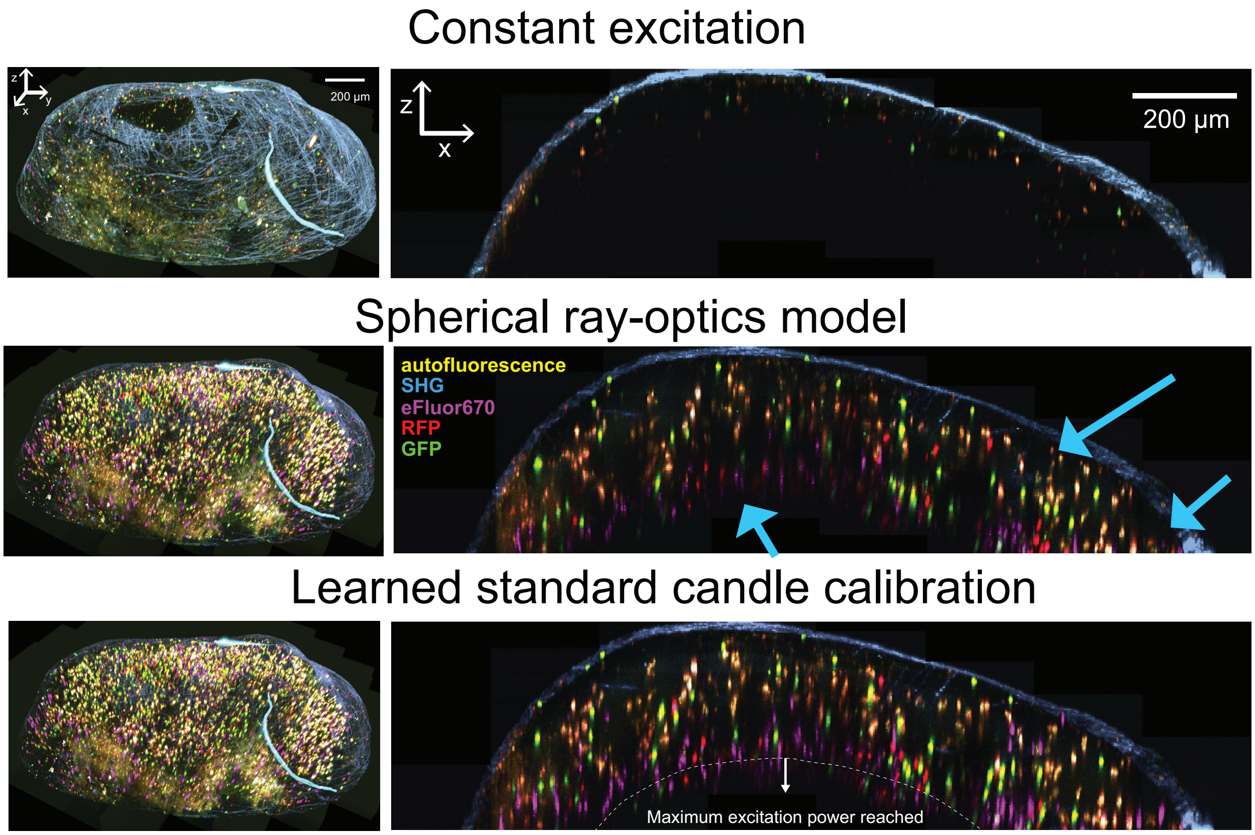


**Figure 4: Comparison of excitation strategies** The same lymph node imaged with constant excitation power (top), spatially varying excitation power as predicted by spherical ray-optics model (middle), or learned standard candle calibration excitation (bottom). Comparing ray optics model excitation and standard candle excitation, several areas can be seen where the former fails to produces low or missing fluorescence from cells that can be clearly seen in the latter (arrows). On the standard candle calibration, a loss of intensity can be seen where the excitation laser reaches maximum power.

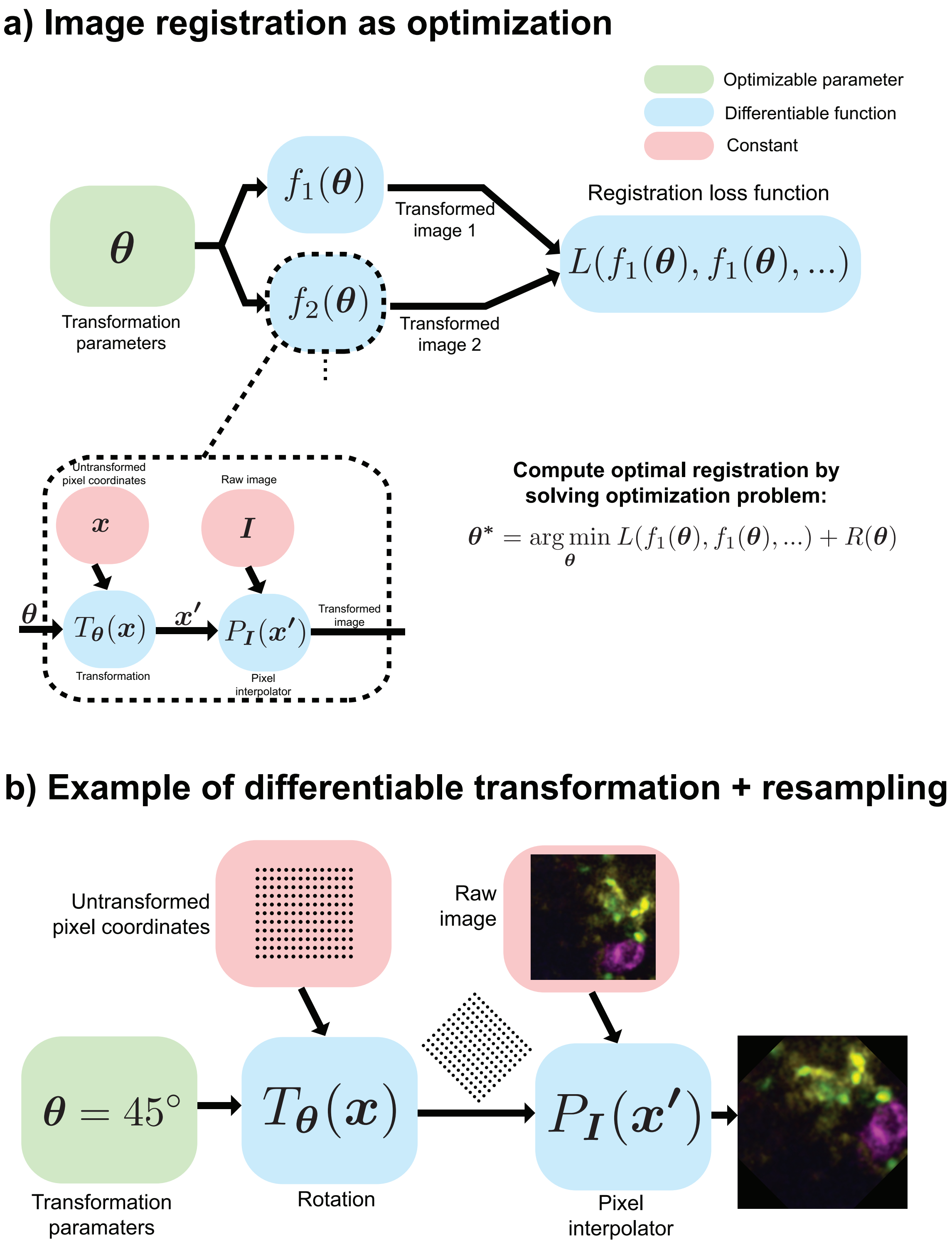


**Figure 5: Image registration formulated as iterative optimization**
a) Overview of *maximum a posteriori* estimation for image registration. Differentiable transformations specific to each correction are used to resample raw image pixels, and fed into a loss function that quantified the quality of solution. b) Rotation as an example of a differentiable transformation.

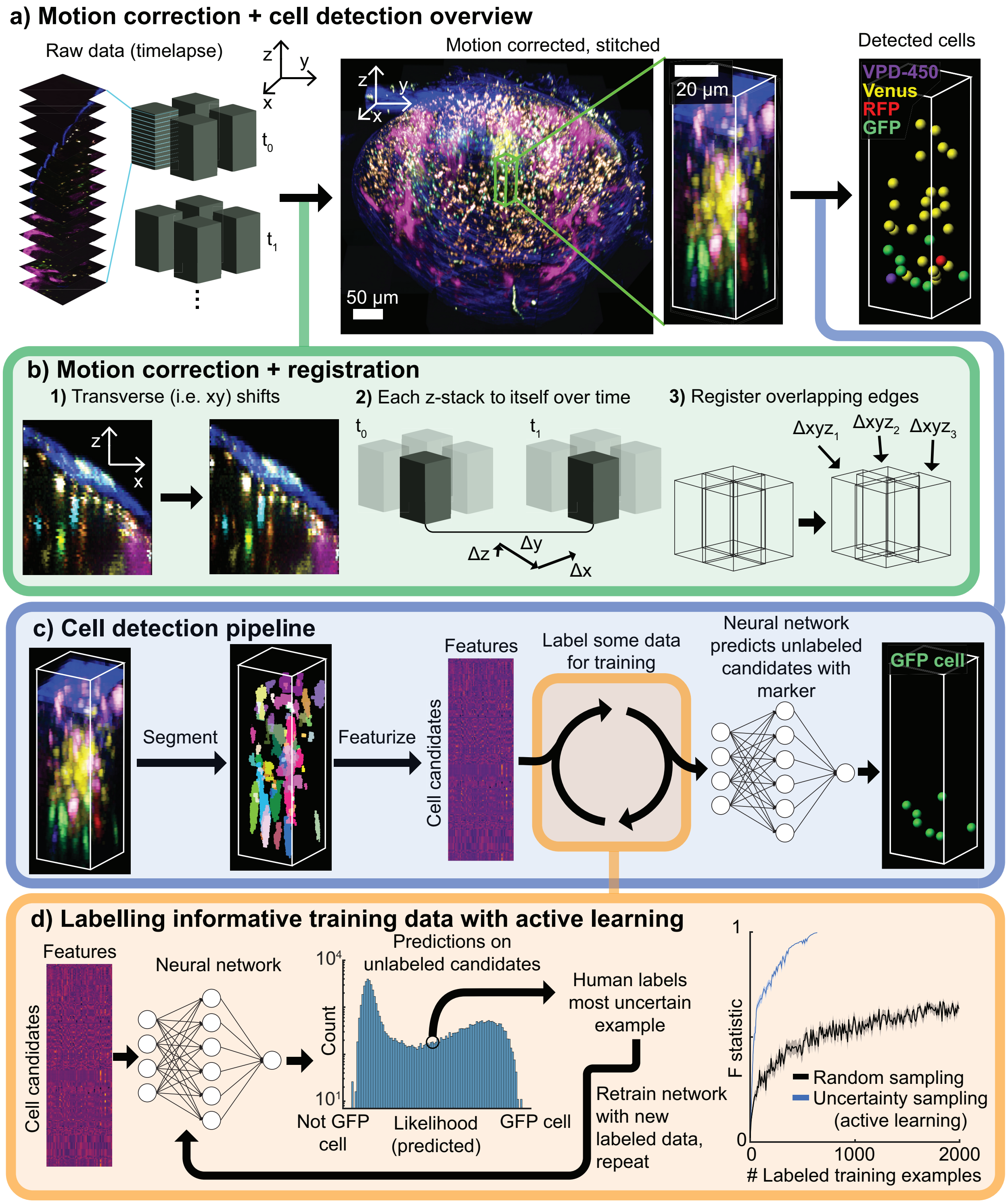


**Figure 6: Motion artifact correction and active learning-based cell detection** a) Overview of data processing converting raw data of separate z-stacks into a single stitched and motion-corrected volume, followed detecting cells based on individual fluorescent protein expression. b) Motion correction and registration consisted of three types of corrections: 1) the XY movements within each slice were optimized. 2) Stacks at consecutive timepoints were registered to one another using cross correlation. 3) The alignment between stacks was optimized. c) Cell identification began by computing a 3D segmentation algorithm to identify candidate cells. Features were then computed for each candidate cell and fed into a classification neural network that predicts which candidates belong to the population of interest d) Active learning was used to label an informative training set. In this paradigm the classification network outputs a measure of certainty that each candidate is a cell or not. The most uncertain of these examples is selected for human labelling, the classification network is retrained, and the procedure is repeated. This enables selection of which candidates belong to population of interest (e.g. GFP). Right, active learning data labelling dramatically boosts classifier accuracy compared to randomly sampling and labelling data.

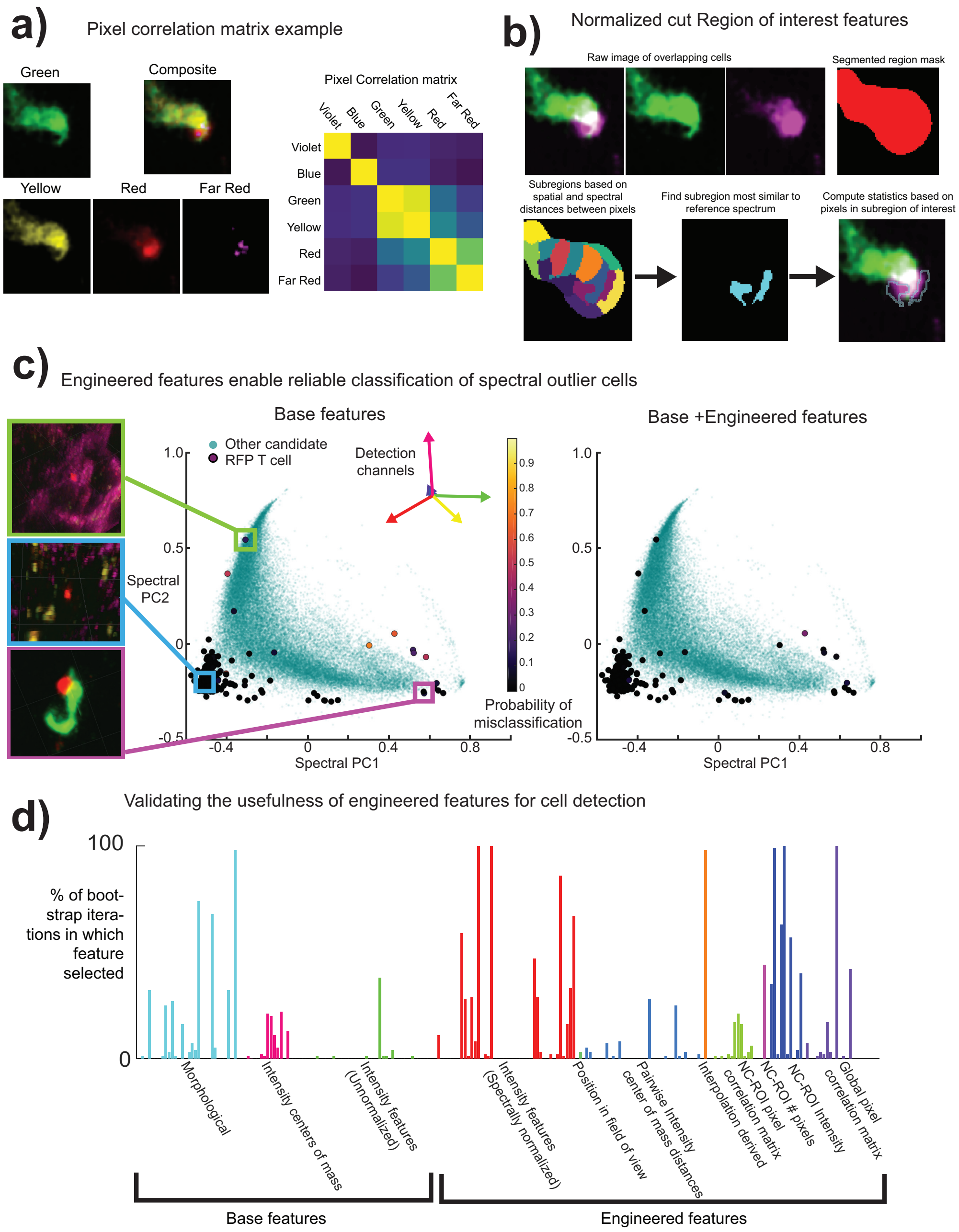


**Figure 7: Engineered features for cell classification** a) The pairwise correlations between pixels of different channels. This provides a clear signal (i.e. distinct clusters in the correlation matrix) when a GFP and RFP labelled cell do not entirely spatially overlap. b) Normalized cut features: by breaking down an area of masked pixels (red, top right) into subregions (bottom left), a subregion that is most similar to a reference spectrum (i.e. the magenta cell) can be identified). c) Including engineered features enables robust identification of spectral outliers. Plots show all candidate cells plotted over first two principal components of the average color spectrum of RFP T cells. Left, spectral outliers (representative images shown on left) from the main cluster of T cells also tend to misclassified. Right, adding in engineered features to classification vastly improves the misclassification probability of these spectral outliers. d) Elastic net bootstrap analysis colored by feature class. Many classes of bootstrapped features were selected a high proportion of the time, validating their usefulness in this classification problem.

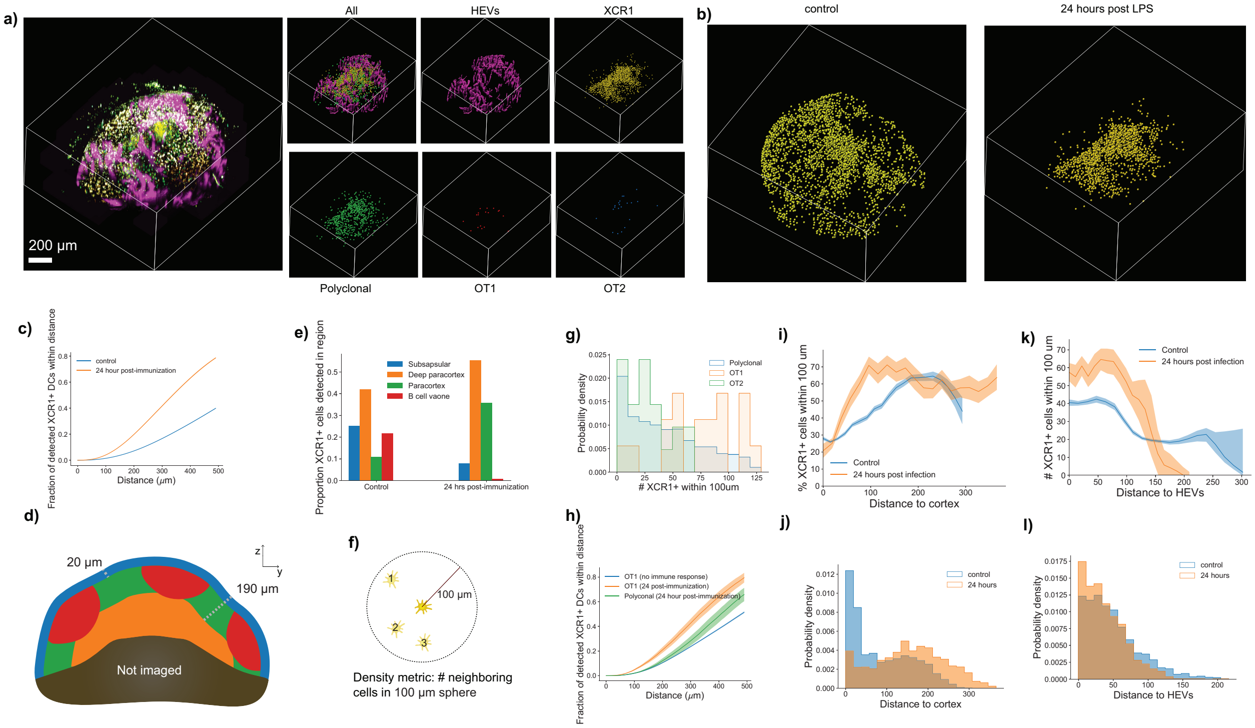


**Figure 8: Reorganization of cell population in the lymph node 24 hours after immunization** a) Image data (left) and localizations of XCR1, Polyclonal, OT1, and OT2 as well as 3D segmentation of high endothelial venules. b) Localization of XCR1 cells in control condition and 24 hours after immunization. c) Amount of clustering as assessed by the mean fraction of XCR1 cells within different distances of XCR1 cells. d) Schematic of how the different parts of the lymph node were defined for e), which shows the changes in localization of XCR1 cells from 0 to 24 hours. f) Schematic of the metric used to assess dendritic cell clustering. g) Histograms of DC cluster density at locations of different types of T cells. h) Fraction of detected XCR1 cells within distance of different types of T cells. Shaded area represents standard error. i) Percent of XCR1 cells within 100 $\mu$m vs. distance to cortex at 0 and 24 hours. Error bars represent bootstrapped 95% confidence interval. j) Histogram of XCR1 cell distances to cortex at 0 and 24 hours. k) Percent of XCR1 cells within 100 $\mu$m vs. distance to HEVs at 0 and 24 hours. i) Histogram of XCR1 cell distances to HEVs at 0 and 24 hours.

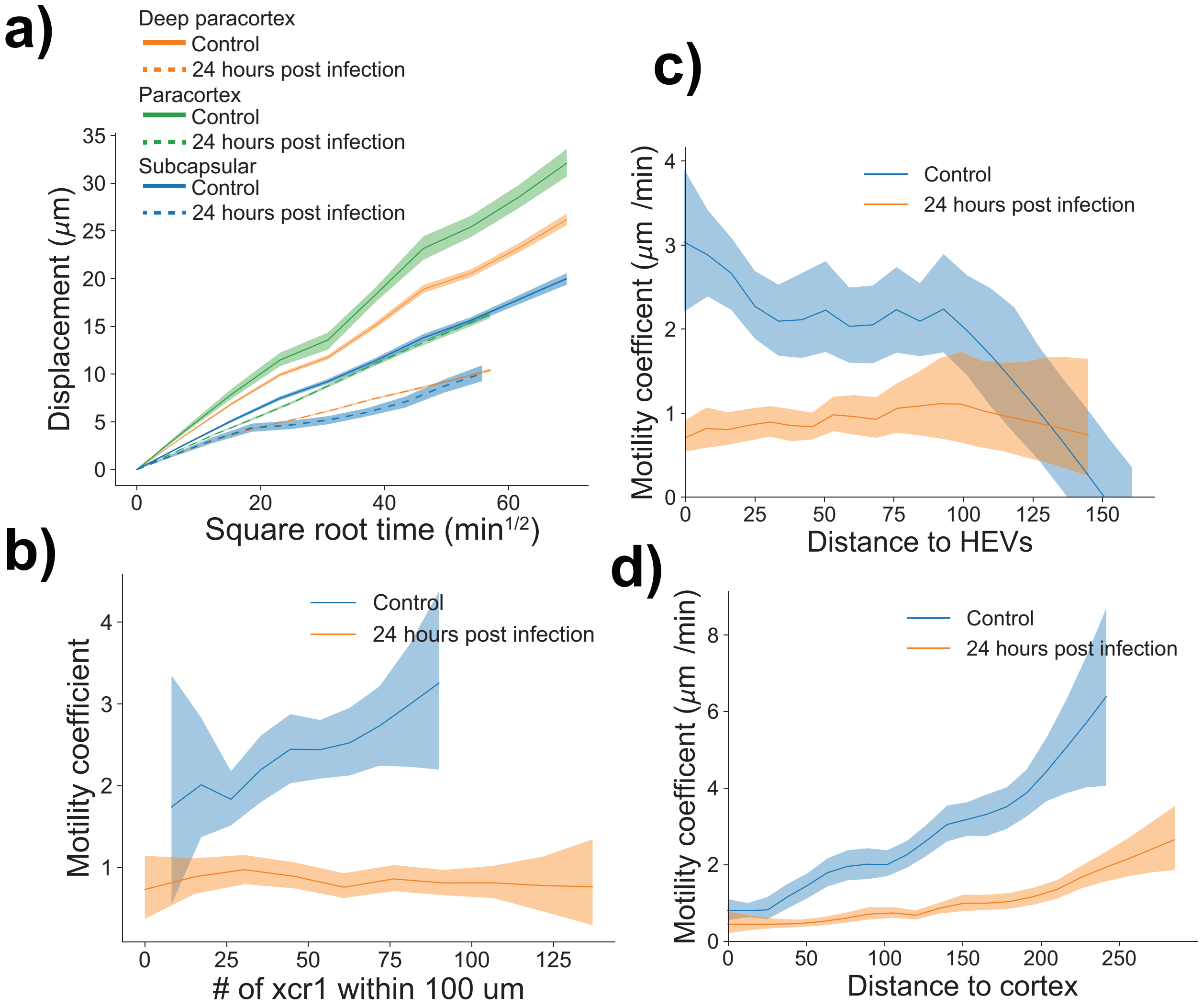


**Figure 9: Dendritic cell motility changes in different anatomical locations** a) Displacement vs. square root time plots for dendritic cells in different parts of the lymph node at 0 and 24 hours. b) Dendritic cell motility coefficients vs the number of other dendritic cells within 100 $\mu$m. c) Motility coefficient vs. distance to high endothelial venules. d) Motility coefficient vs. distance to cortex. Shaded regions in all plots represent bootstrapped 95% confidence intervals.
